# Gut microbiome derived folate metabolite suppresses colorectal cancer progression

**DOI:** 10.64898/2026.07.14.738490

**Authors:** Rebecca Danner, Juyeong Cho, Zachary Detwiler, Jonathan Williams, Jiyoung Amanda Han, Chang Yang, Xavier Diebold, Katerina Maeder, Jonathan G. Van Vranken, Allison S. Walker, Cammie Lesser, Snehal N. Chaudhari

**Author notes:** Corresponding author: Snehal N. Chaudhari PhD University of Wisconsin-Madison, 433 Babcock Drive, Madison, WI 53706.

## Abstract

The gut microbiota influences colorectal cancer (CRC) progression, primarily through the secretion of small molecule metabolites. While numerous microbial products are known to drive CRC, endogenous protective mechanisms remain largely uncharacterized. Utilizing a folate metabolomics platform, we demonstrate that the healthy gut microbiota produces folinic acid (FA), a known chemotherapeutic adjuvant also known as leucovorin. This microbially derived folinic acid is progressively depleted in mouse models of colitis-associated CRC and in human clinical metagenomic cohorts with advancing disease severity. Mechanistically, folinic acid acts as a signaling molecule that directly binds and inhibits the intracellular protease calpain-2. This interaction stabilizes epithelial E-cadherin protein expression and suppresses CRC epithelial-to-mesenchymal transition driving metastasis. Genetically manipulating gut microbial production of FA is sufficient to modulate CRC *in vivo*, even in the presence of chronic inflammation. This study reframes folinic acid from a chemotherapeutic enhancer to an endogenous microbial metabolite that actively suppresses CRC progression.

## Main

Leucovorin, or folinic acid, is a naturally occurring folate metabolite widely used in chemotherapy regimens designed to treat advanced stages of colorectal cancer (CRC)^1,2^. A combinatorial treatment regimen of folinic acid and 5-flurouracil, often with other cytotoxic agents, is the current standard of care for neoadjuvant, adjuvant, and palliative therapy for CRC^3^. Folinic acid is known to enhance cytotoxic effects of 5-flurouracil by forming a ternary complex that stabilizes and inhibits thymidylate synthase, an enzyme crucial for DNA synthesis and repair^4^. Pharmacologically, folinic acid is versatile as it is used both to enhance 5-flurouracil cytotoxicity and to counteract the cytotoxic effects of antifolate chemotherapeutics such as methotrexate^5^. Both applications of folinic acid require participation in the one-carbon metabolism cycle. Unlike folic acid, folinic acid is naturally occurring and biologically active as it does not require dihydrofolate reductase for conversion into tetrahydrofolate for entry into the folate cycle^6^. However, recent evidence suggests that folinic acid can also function as a signaling molecule in cells, potentiating the cytotoxicity of non-fluoropyrimidine agents such as the proteasome inhibitor bortezomib through the upregulation of apoptotic signaling^7^. This suggests a multifaceted anti-cancer role for folinic acid, with yet unknown therapeutic potential that extends beyond the classic ternary complex with thymidylate synthase. Uncovering these signaling pathways may revolutionize therapy for CRC, which remains the second leading cause of cancer mortality worldwide despite advancements in screening and diagnosis^8^.

As an essential component of the colon, and thus of the CRC tumor microenvironment, the gut microbiota has been associated with CRC progression^9,10^. A healthy gut microbiome plays a critical role in the maintenance of gut homeostasis by supporting digestion, producing metabolites, and regulating the immune system^11^. When this microbial balance is disrupted, a dysbiotic microbiota can contribute to CRC progression by promoting chronic inflammation, increasing immune dysregulation, and altering metabolism^10^. It is widely known that CRC is characterized by a decrease in abundance of health-associated bacteria, alongside a concomitant enrichment of pathogenic bacteria^12,13^. Recent studies have highlighted that the direct influence of the gut microbiome on colorectal cancer is mediated predominantly by bacterial metabolites^14,15^. Gut microbes produce metabolites that can either protect against or promote cancer. Harmful metabolites such as secondary bile acids, sulfides, and other toxins can damage DNA, trigger inflammation, and promote tumor development^11^. In contrast, beneficial metabolites like the short-chain fatty acids butyrate, acetate, and propionate promote anti-inflammatory and anti-cancer effects by supporting intestinal health and activating tumor-suppressing genes^16^. Whether gut microbial folate metabolites influence CRC development is unknown.

To investigate the host-microbe folate axis, we recently performed a systematic analysis of folate metabolites in 37 tissues from male and female mice under germ-free and conventional conditions^17^. We found that a healthy gut microbiota consumes folate, with the germ-free gastrointestinal tract showing the largest increase in folate availability compared to conventional among the 37 tissues analyzed. Considering that excess folate is shunted into intracellular storage as folinic acid^18^, we found that folinic acid accumulated in the lower gastrointestinal tracts of colonized mice and was absent in germ-free counterparts. These data identify the gut microbiota as the biosynthesis source of the intestinal folinic acid pool. Given the clinical importance of folinic acid in modulating CRC chemotherapy, its microbial origin represents a previously unrecognized endogenous defense mechanism against CRC progression.

In this study, we find that folinic acid production by the gut microbiome is reduced in CRC mouse models and human patients, particularly in patients with metastatic CRC. We show that microbial folinic acid enhances gut epithelial integrity and suppresses CRC progression. This protective efficacy of folinic acid is independent of its ability to mitigate gut inflammation, providing anti-neoplastic benefits even in the presence of chronic colitis. Investigation into the molecular mechanisms identify folinic acid as an inhibitor of calpain-2 (CAPN2), an intracellular protease known to promote epithelial-to-mesenchymal transition of CRC cells. We demonstrate that genetically manipulating gut microbial folinic acid production is sufficient to influence CRC progression. Thus, this study identifies an unexplored metabolic checkpoint, positioning the gut microbiome-folate axis as a key determinant of CRC.

## Results

### Bacterial folinic acid production is decreased in murine and human colorectal cancer

In the one-carbon metabolism cycle, folates serve as essential cofactors facilitating the transfer of one-carbon units for key anabolic processes, including DNA synthesis, repair, and methylation^19^ (Figure 1a). Folinic acid (5-formyltetrahydrofolate) is not a one-carbon donor, rather it is a chemically stable reservoir of reduced folates produced from 5,10-methylenetetrahydrofolate via a side reaction catalyzed by the serine hydroxymethyltransferase (SHMT) enzyme, which in bacteria is encoded by the *glyA* gene^20^ (Figure 1a). Utilizing our established folate liquid chromatography-mass spectrometry (LC-MS) metabolomics platform^17^, we verified that intestinal folinic acid levels are dependent on the presence of a functioning microbiota. Folinic acid was detected in the gut of conventionally colonized mice but was virtually absent in germ-free mice (Figure 1b). These data establish the intestinal microbiota as the primary source of luminal folinic acid under folate-replete conditions. To evaluate the relevance of microbially-produced folinic acid in the context of colorectal cancer (CRC), we extended our analysis to the C57BL/6J-*Il10^-/-^*(IL10KO) mouse model of colitis-associated CRC^21^. Strikingly, cecal abundance of folinic acid was ∼10-fold lower in IL10KO mice compared to wild-type (WT), suggesting that microbiota in IL10KO mice are deficient in folinic acid production (Figure 1c).

**Fig. 1.**
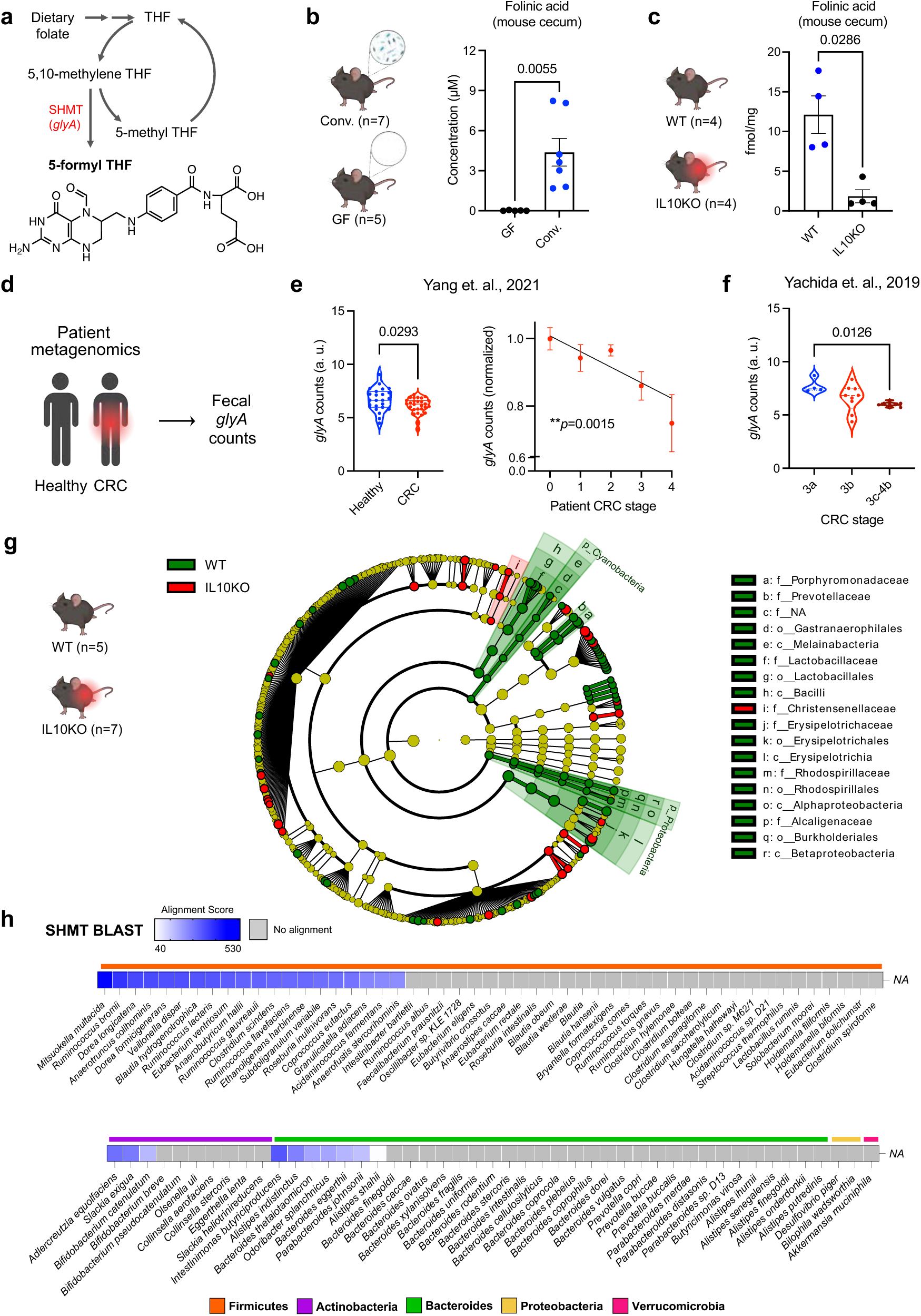
Bacterial folinic acid production is decreased in murine and human colorectal cancer. **a**. Schematic of one-carbon metabolism that produces folinic acid (5-formyl THF). **b**. Folinic acid concentration detected in the cecum of germ-free (GF) and conventionally colonized (Conv.) mice via LC-MS. (n=5 GF and n=7 Conv.; two-tailed Welch’s t test). **c**. Concentration of folinic acid in the cecum of wild-type (C57BL/6) and colitis-associated CRC (IL10KO) mice. (n=4 per group; Mann-Whitney test). **d-f**. *glyA* gene counts in fecal metagenomic datasets from patients with colorectal cancer (CRC) normalized to people without colorectal cancer at various stages of disease (Yang et. al., healthy, n=23; CRC, n=28, Mann-Whitney test; Yachida et. al., 3a, n=4; 3b, n=9; 3c-4b, n=9, One-way ANOVA Krustal-Wallis test). **g.** LEfSe (Linear discriminant analysis Effect Size) cladogram analysis of microbiota. (n=5 WT and n=7 IL10KO). **h**. Heatmap of SHMT BLAST alignment scores across 119 representative gut bacterial species spanning five phyla. Grey represents no alignment. All statistically significant *p* values are indicated in each bar graph. All bar graphs are represented as mean ± SEM, data points represent biological replicates.

To determine whether a loss of microbial folinic acid production is also a hallmark of human CRC, we assessed *glyA* gene abundance in CRC patient microbiota (Figure 1d). Metagenomic analysis of human stool samples revealed a significant reduction in *glyA* gene counts in patients with CRC compared to healthy controls^22^ (Figure 1e). Strikingly, we observed a progressive decrease in microbial *glyA* abundance tracking with advanced disease severity, suggesting that depletion in microbial folinic acid production is coupled to CRC malignancy and metastasis (Figure 1e,f)^22,23^.

To investigate the bacterial drivers behind the observed depletion of fecal folinic acid in CRC, we performed microbiota 16S rRNA sequencing in fecal samples from IL10KO mice and WT controls. Linear Discriminant Analysis Effect Size (LEfSe) cladogram analysis revealed an inflammatory and pro-CRC signature in IL10KO animals, characterized by an expansion of *Christensenellaceae* family, and a loss of homeostatic clades such as *Lactobacillaceae, Bacilli,* and several lineages within *Proteobacteria* (such as *Burkholderiales*, *Alphaproteobacteria*, and *Betaproteobacteria*) (Figure 1g). Abundant anaerobic classes like *Clostridia* and *Bacteroidia* did not display uniform, class-wide shifts. To determine if these shifts corresponded to a loss of folinic acid biosynthetic capacity, we performed a BLAST analysis of the bacterial SHMT protein encoded by the *glyA* gene across the defined 119-member human commensal (hCom2) community^24^ to identify the taxonomic distribution of *glyA.* This analysis revealed that *glyA* is conserved within specific obligate anaerobes clustering within groups of core homeostatic gut commensals, including dominant lactic acid bacteria and major carbohydrate-degrading *Clostridiales* lineages, which are depleted in CRC^12,13^ (Figure 1h). The SHMT-positive *Lactobacillaceae* and *Proteobacteria* lineages that dominate the folinic acid-replete WT gut were also significantly depleted in the IL10KO mice (Figure 1g,h). Taken together, these data support a model wherein the fecal folinic acid deficiency observed in CRC is driven not by broad shifts in abundant bacterial taxa, rather by the targeted depletion of specific *glyA*-containing bacterial strains.

### Folinic acid increases E-cadherin expression and suppresses epithelial-to-mesenchymal transition in human CRC cells

Folinic acid is considered a non-toxic folate vitamer and acts as a mucosal protective agent in clinical settings^25^. We confirmed that folinic acid does not, by itself, kill cancer cells, by incubating human colorectal adenocarcinoma epithelial HT29 cells with varying concentrations of folinic acid for 4 days *in vitro* (Figure 2a). While no overt toxic effects were observed in a cell viability assay, HT29 cells incubated with folinic acid exhibited a dramatic morphological restructuring of colonies. The vehicle-treated controls behaved like HT29 cells in culture; rather than arresting to form a structured epithelial monolayer upon confluence, these cells stacked into overlapping, mesenchymal-like disorganized multilayers, characteristic of undifferentiated parental HT29 cells^26^ (Figure 2b). However, HT29 cells treated with folinic acid appeared more epithelial and less mesenchymal in morphology. Volume/area quantification of each colony confirmed that folinic acid induced a monolayer-like re-structuring of cells compared to controls (Figure 2b). Based on this observation, we hypothesized that folinic acid prevents or reverses epithelial-to-mesenchymal transition (EMT) of CRC cells *in vitro* (Figure 2c). Loss of epithelial cell-to-cell adherens junction proteins, such as E-cadherin, is a hallmark of EMT in CRC^27^ (Figure 2c)Click or tap here to enter text.. Consistently, E-cadherin levels were significantly increased in HT29 cells treated with folinic acid compared to the vehicle treated cells as measured via immunocytochemistry (Figure 2d) and western blot (Figure 2e) analysis. Thus, while folinic acid is not cytotoxic, it increases E-cadherin protein levels in CRC cells *in vitro*.

**Fig. 2.**
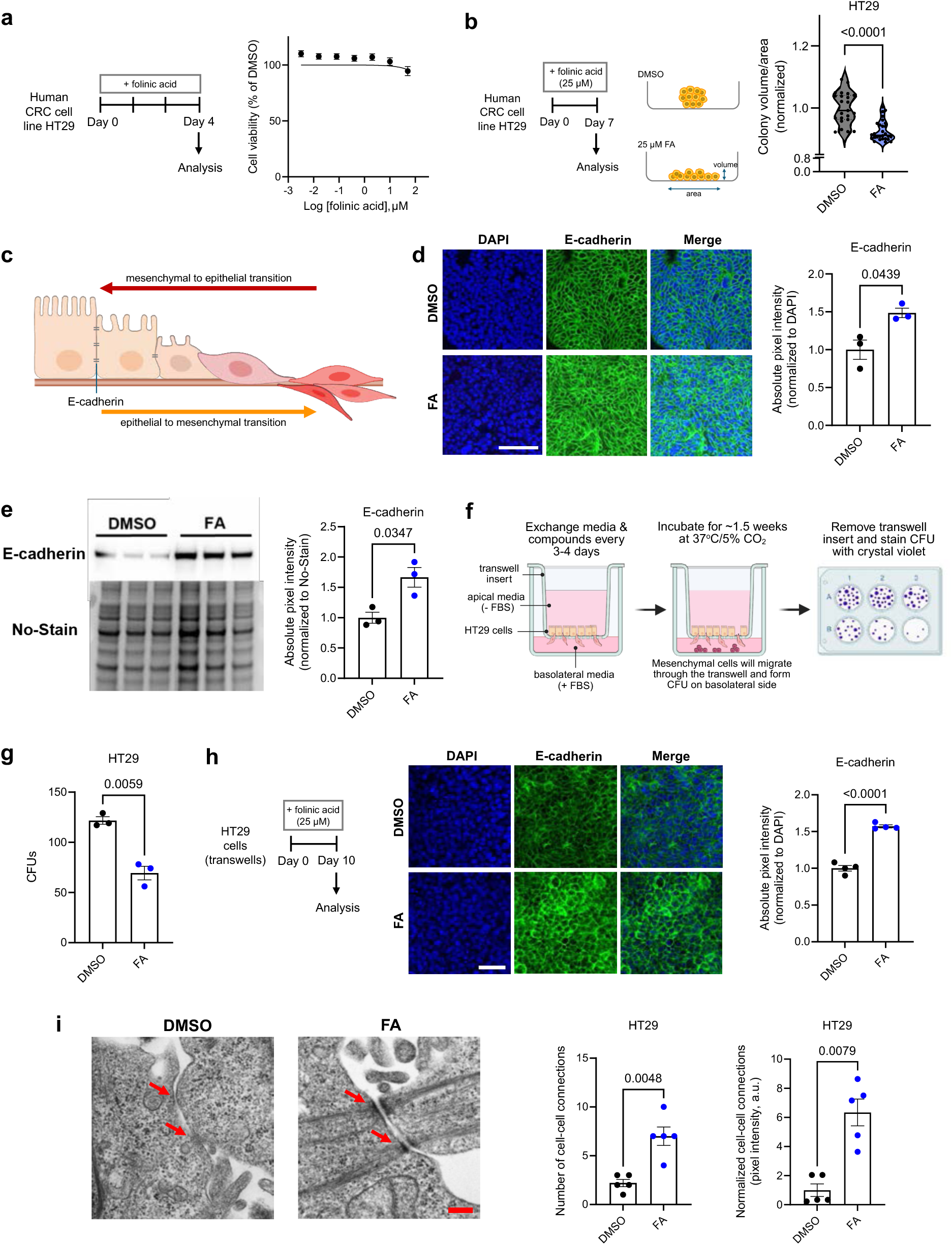
Folinic acid increases E-cadherin expression and suppresses epithelial-to-mesenchymal transition in human CRC cells. **a**. Cell viability of HT29 cells treated with folinic acid (0.0032, 0.016, 0.08, 0.4, 2.0, 10, and 50 µM) for 96 hours, as measured by CellTiter-Glo assay. **b**. Colony volume/area of each colony forming unit (CFU) from HT29 cells treated with 25 µM folinic acid (FA) or 0.5% DMSO for 7 days (n=23 DMSO and n=28 FA.; Mann-Whitney test). **c**. Schematic of epithelial-to-mesenchymal transition (EMT). **d**. Representative images of HT29 cells stained with E-cadherin and DAPI. Scale bar, 100 µm. Absolute pixel intensity of E-cadherin was measured using ImageJ and normalized to DAPI. (n=3 per group, two-tailed Welch’s t test). **e**. Western blot of E-cadherin expression normalized to No-Stain (n=3 per group, two-tailed Welch’s t test). **f**. Schematic of transwell assay to analyze EMT. **g**. Colony forming units (CFUs) of HT29 cells treated with 0.5% DMSO or 25 µM folinic acid (FA) (n=3 per group, two-tailed Welch’s t test). **h**. Representative images of apical HT29 cells stained with E-cadherin and DAPI. Scale bar, 100 µm. Absolute pixel intensity of E-cadherin was measured using ImageJ and normalized to DAPI. (n=4 per group, two-tailed Welch’s t test). **i**. Representative transmission electron microscopy images of HT29 cells treated with 0.5% DMSO or 25 µM folinic acid (FA). Scale bar, 200 nm. Red arrows indicate cell-cell connections (n=5 per group, two-tailed Welch’s t test for number of cell-cell connections and Mann-Whitney test for pixel intensity). All *p* values are indicated in each bar graph. All bar graphs are represented as mean ± SEM, data points represent biological replicates.

Next, we wanted to investigate whether an increase in E-cadherin and enhancement of epithelial-like morphology in HT29 cells is sufficient to reduce its metastatic capacity *in vitro*. Using a transwell migration assay^28^ (Figure 2f), we found that HT29 cells treated with folinic acid were less migratory as determined by the significantly decreased number of colony forming units (CFUs) on the basolateral side (Figure 2g). Consistent with our previous results, folinic acid-treated cells in the transwells showed high E-cadherin levels (Figure 2h). Transmission electron microscopy analysis of cells in transwells also displayed an increase in the number and protein density of cell-cell connections in the folinic acid treatment compared to the controls (Figure 2i). These data demonstrate that folinic acid regulates CRC cell plasticity by maintaining E-cadherin expression, reinforcing epithelial cell-to-cell junctions, and directly reducing migratory capacity *in vitro*.

### Folinic acid inhibits calpain-2 and reverses CRC epithelial-to-mesenchymal transition

We next sought to identify the mechanism of how folinic acid reduces EMT in CRC cells. Using target-engagement proteomic analysis on HT29 cell lysates incubated with folinic acid, we identified calpain-2 (CAPN2) and its catalytic subunit (CAPNS) as potential binding targets (Figure 3a). This interaction was further supported by *in silico* docking analysis, which showed that folinic acid binds to the catalytic subunit of CAPN2 (Figure 3b). Notably, the 5-formyl group of the bound folinic acid is oriented toward the catalytic cysteine in the CAPN2 active site, suggesting that this folate vitamer acts as a selective inhibitor of calpain protease activity. To biochemically characterize this ligand-protein interaction, we cloned, overexpressed, and purified human CAPNS for *in vitro* binding assays. Recombinant protein was incubated with equimolar concentrations of five distinct folate vitamers, and protein-bound folates were subsequently quantified via LC-MS to evaluate competitive binding affinity. Folinic acid (5-formyl-tetrahydrofolate) exhibited the highest binding affinity to the CAPN2 catalytic subunit, followed by 5-methyl-tetrahydrofolate (Figure 3c). This pattern indicates that modifications at the *N*^5^ position of the pterin ring are critical determinants of CAPN2 binding. To test whether folinic acid functions as a competitive inhibitor, we performed a ligand displacement assay using a known, potent covalent inhibitor of CAPN2 (CAPN2i, CAS No: 110115-07-6). Purified protein pre-incubated with folinic acid demonstrated dose-dependent displacement of the vitamer upon the addition of CAPN2i as measured by LC-MS (Figure 3d). These data confirm that folinic acid and CAPN2i competitively occupy the same catalytic binding pocket.

**Fig. 3.**
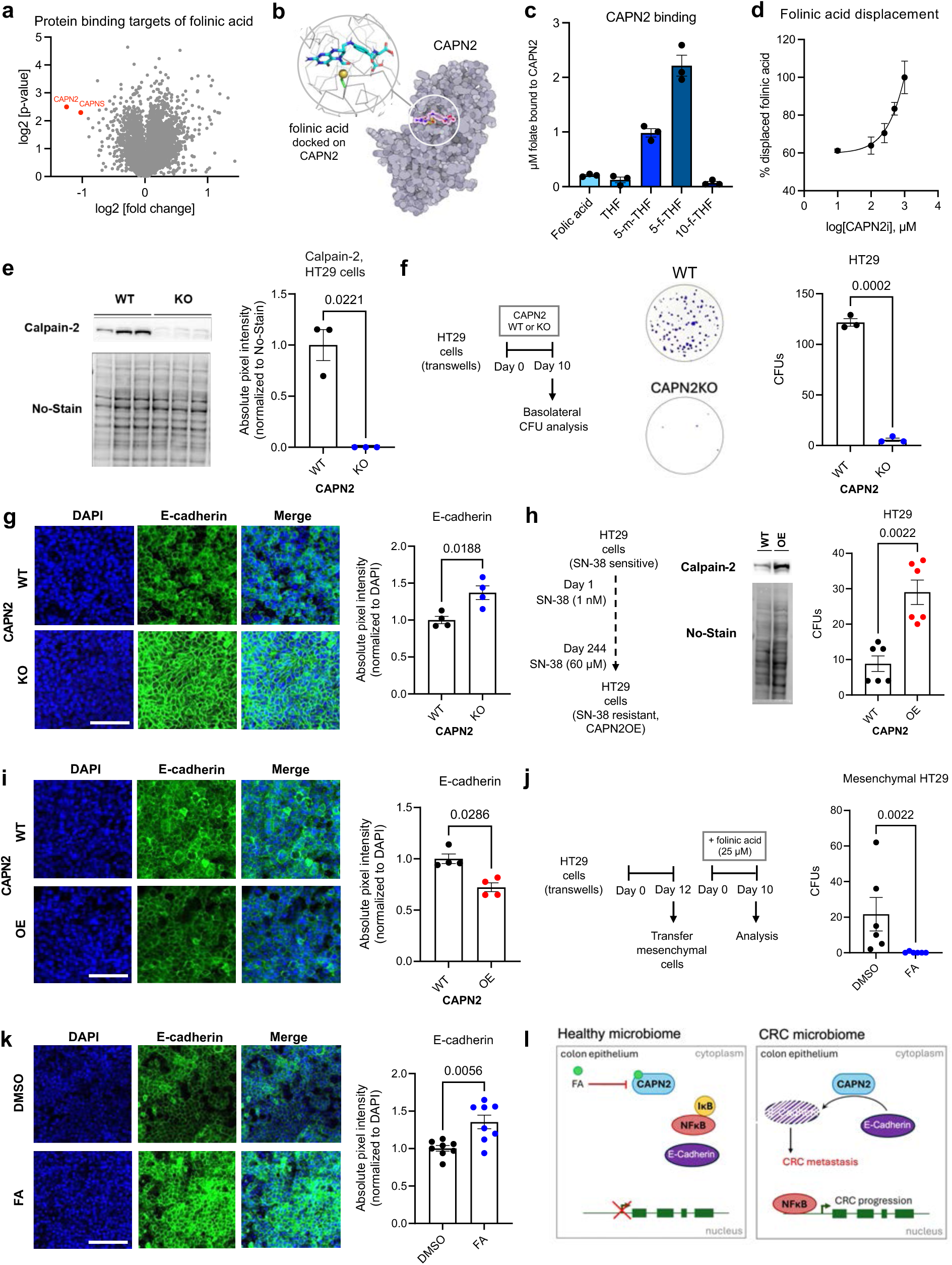
Folinic acid inhibits calpain-2 and reverses CRC epithelial-to-mesenchymal transition. **a**. Target-engagement proteomic analysis of HT29 cell lysates incubated with folinic acid showing calpain-2 (CAPN2) and its catalytic subunit (CAPNS) as significant binding targets indicated in red. **b**. *In-silico* docking of folinic acid and CAPN2, with the catalytic cysteine shown in insert. **c**. *In vitro* recombinant CAPNS binding assay using LC-MS for folic acid, tetrahydrofolate (THF), 5-methyl-tetrahydrofolate (5-m-THF), 5-formyl-tetrahydrofolate (5-f-THF or folinic acid), and 10-formyl-tetrahydrofolate (10-f-THF). **d**. Folinic acid displacement assay using CAPN2i measured via LC-MS. (n=3 per concentration). **e**. Western blot of CAPN2 expression normalized to No-Stain in wild-type (WT) and CAPN2 knockout (CAPN2KO) HT29 cells (n=3 per group, two-tailed Welch’s t test). **f**. Colony forming units (CFUs) of WT and CAPN2KO HT29 cells from transwell assay. (n=3 per group, two-tailed Welch’s t test). **g**. Representative images of apical HT29 cells stained with E-cadherin and DAPI. Scale bar, 100 µm. Absolute pixel intensity of E-cadherin was measured using ImageJ and normalized to DAPI. (n=4 per group, two-tailed Welch’s t test). **h**. Western blot of CAPN2 expression normalized to No-Stain in wild-type (WT) and CAPN2 overexpressed (CAPN2OE) HT29 cells. CFUs of WT and CAPN2OE HT29 cells from transwell assays. (n=6 per group, Mann-Whitney test). **i**. Representative images of apical HT29 cells stained with E-cadherin and DAPI. Scale bar, 100 µm. Absolute pixel intensity of E-cadherin measured using ImageJ and normalized to DAPI. (n=4 per group, Mann-Whitney test). **j.** CFUs of mesenchymal HT29 cells treated with 0.5% DMSO or 25 µM folinic acid (FA) (n=6 per group, two-tailed Welch’s t test). **k**. Representative images of apical HT29 cells stained with E-cadherin and DAPI. Scale bar, 100 µm. Absolute pixel intensity of E-cadherin measured using ImageJ and normalized to DAPI. (n=8 per group, two-tailed Welch’s t test). **l**. Model of how microbial folinic acid (FA) inhibits CAPN2 to reduce CRC progression. All *p* values are indicated in each bar graph. All bar graphs are represented as mean ± SEM, data points represent biological replicates.

Previous studies have established that CAPN2 is overexpressed in CRC and inhibiting its activity suppresses tumor burden *in vivo*^29,30^. Mechanistically, CAPN2 is known to promote CRC progression by driving a pro-inflammatory gene signature, a process mediated by activation of the pro-inflammatory transcription factor NF-κB^29^. Given our discovery that folinic acid inhibits CAPN2 and suppresses EMT *in vitro*, we next sought to investigate whether CAPN2 natively promotes EMT in CRC cells and directly modulates expression of the epithelial marker E-cadherin. To evaluate this, we utilized CRISPR-Cas9 genome editing to generate a CAPN2 knockout (KO) HT29 cell line (Figure 3e) and performed transwell migration assays alongside E-cadherin quantification. While the WT HT29 cells exhibited robust migratory capacity, CAPN2KO cells displayed a striking ∼10-fold reduction in migration, suggesting that depletion of CAPN2 is sufficient to attenuate the EMT phenotype of CRC cells *in vitro* (Figure 3f). Consistently, CAPN2KO cells demonstrated markedly elevated E-cadherin expression compared to WT controls (Figure 3g).

Given that CAPN2 is frequently overexpressed in aggressive and chemotherapy-resistant CRC tumors^31,32^, we next investigated whether elevated CAPN2 expression is independently capable of driving EMT. To test this, we generated a CAPN2-overexpressing (OE) cell line by progressively treating HT29 cells with increasing doses of SN-38, the active metabolite of the chemotherapy drug irinotecan^29^ (Figure 3h)Click or tap here to enter text.. These SN-38-resistant cells exhibited a marked upregulation of CAPN2, which coincided with a significant increase in cell migration and a concomitant decrease in E-cadherin expression (Figure 3h,i). These data indicate that elevated CAPN2 protein levels are a hallmark of chemoresistance-associated EMT in CRC.

Finally, to determine whether folinic acid treatment can actively reverse an established mesenchymal phenotype, we isolated the migrated, mesenchymal-like CRC cells from the bottom of the initial transwell chambers and transferred them to secondary transwell assays for treatment with folinic acid (Figure 3j). Strikingly, folinic acid administration potently suppressed the subsequent migration of these pre-isolated mesenchymal cells. This phenotypic rescue was supported by a concomitant upregulation of E-cadherin protein levels (Figure 3j,k), altogether demonstrating that folinic acid inhibits CAPN2, which can reduce CRC progression and reversing EMT (Figure 3l).

### Gut microbial synthesis of folinic acid improves colon epithelial and mucosal homeostasis in vivo

To determine if microbial folinic acid promotes E-cadherin expression and intestinal homeostasis *in vivo*, we utilized the probiotic commensal *Escherichia coli* Nissle 1917 (*EcN*), which can naturally produce all folate derivatives^33^. Using Lambda Red recombination^34^, we deleted the *glyA* gene to generate a genetically engineered *EcN* strain (*EcN glyA* KO) with a dampened capacity of producing folinic acid (Extended Data Figure 1). C57BL/6J mice were gavaged biweekly with 10^9^ CFUs of either wild-type (WT) *EcN* or *EcN glyA* KO for 12 weeks (Figure 4a). While both strains colonized at equivalent levels (Figure 4b), LC-MS analysis confirmed a significant reduction in fecal folinic acid production in the *glyA* KO group (Figure 4c).

**Fig. 4.**
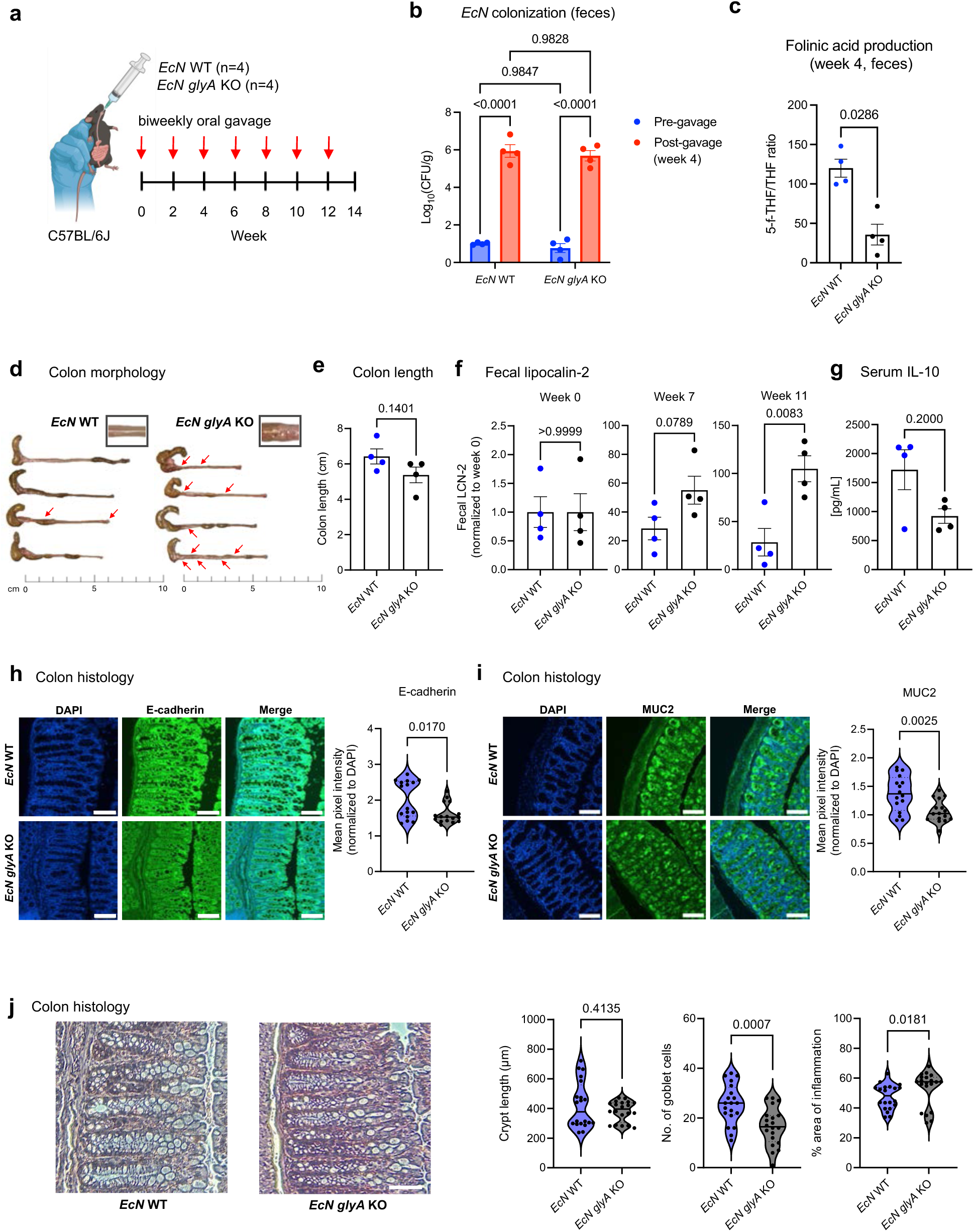
Microbial synthesis of folinic acid improves epithelial and mucosal homeostasis in C57BL/6J colon. **a.** Schematic of a biweekly oral gavage of *E. coli* Nissle wild-type (WT) or *glyA* KO (10^9^ CFUs) administered to C57BL/6J mice at 6 weeks of age, followed by colon collection after 14 weeks. **b.** *EcN* WT and *EcN glyA* KO colonization by RT-qPCR in mouse feces **c.** Fecal folinic acid levels measured by LC-MS. **d.** Colon images with irregularities marked with red arrows. **e.** Colon length (cm) of each mouse. (n=4 per group, two-tailed Welch’s t test). **f.** Fecal lipocalin-2 (LCN-2) levels measured by ELISA in samples collected at week 0, 7 and 11. Each n represents one mouse (n=4 per group, two-tailed Welch’s t test). **g.** IL10 cytokine level in mouse serum (n=4 per group, two-tailed Mann Whitney test). **h.** Representative images of mouse colons stained with E-cadherin and DAPI. Scale bar, 100 µm. Mean pixel intensity of E-cadherin was measured from 4 images per mouse colon using ImageJ and normalized to DAPI. (n=16 per group, Mann-Whitney test). **i.** Representative images of mouse colons stained with mucin 2 (MUC2) and DAPI. Scale bar, 100 µm. Mean pixel intensity of MUC2 was measured from 4 images per mouse colon using ImageJ and normalized to DAPI. (n=16 per group, two-tailed Welch’s t test). **j.** Representative images of mouse colons stained with hematoxylin and eosin (H&E). Scale bar, 100 µm. All images were analyzed in a blinded manner. Crypt length (µm) and number of goblet cells were measured by QuPath and the % area of inflammation was measured by ImageJ based on 5 crypts per mouse. Each n represents one crypt. (n=20 per group; two-tailed Welch’s t test (goblet cells) and Mann-Whitney test (crypt length and % area of inflammation)). All *p* values are indicated in each bar graph. All bar graphs are represented as mean ± SEM, data points represent biological replicates.

After 14-weeks, animals were sacrificed and colon tissue was harvested for analysis. Strikingly, the *EcN glyA* KO colonized mouse colons appeared to exhibit signs of inflammation, characterized by visible thickening and increased irregularities along the length of the colon compared to the *EcN* WT group^35^ (Figure 4d). This morphological change occurred with a slight but not significant reduction in colon length in *EcN glyA* KO gavaged mice compared to the *EcN* WT mice, which is indicative of intestinal inflammation and tissue damage^36^ (Figure 4e). Consistent with these macroscopic changes, fecal levels of lipocalin-2 (LCN-2), a biomarker of intestinal inflammation^37^, showed a progressive and significant increase by week 11 in the *glyA* KO group (Figure 4f). Consistently, the serum levels of the anti-inflammatory cytokine interleukin 10 (IL-10)^38^, was decreased in the *EcN glyA* KO gavaged mice compared to the WT (Figure 4g). Furthermore, immunofluorescent staining of colon sections showed that the WT-colonized mice maintained improved epithelial and mucosal homeostasis, evidenced by significantly higher E-cadherin (Figure 4h) and mucin 2 (MUC2) (Figure 4i) expression compared to the *EcN glyA* KO group. Hematoxylin and eosin (H&E) staining and histological analysis of colon sections further revealed that the *glyA* KO colonization was associated with increased inflammatory infiltration and a depletion of goblet cells, despite no significant difference in crypt length (Figure 4j). Collectively, these data suggest that microbially derived folinic acid is essential for maintaining intestinal homeostasis and preventing inflammation.

### Microbial synthesis of folinic acid suppresses CRC progression in the presence of chronic inflammation *in vivo*

Based on our results in wild-type C57BL/6J mice, we next analyzed the effect of bacterial folinic acid production in increasing E-cadherin expression and reducing chronic inflammation in the colitis-associated C57BL/6J-*Il10^-/-^* (IL10KO) mouse model. IL10KO mice were gavaged biweekly with either *EcN* WT or *EcN glyA* KO for 20 weeks (Extended Data Figure 2a). After 21-weeks, animals were sacrificed, and colon tissue was harvested for analysis. As expected, all IL10KO mouse colons exhibited significant signs of colitis, including cecal blood and structural thickening, compared to wild-type controls (Extended Data Figure 2b). While the *EcN* WT colonized colons exhibited a more uniform, less thickened morphology compared to the *EcN glyA* KO group, the effect did not appear to be consistent across all mice. No significant change in colon length was observed between groups (Extended Data Figure 2c). Consistently, while *EcN* WT colonization was associated with a significant reduction in fecal LCN-2 at week 10, this anti-inflammatory effect was abolished by week 21 (Extended Data Figure 2d). Immunofluorescent analysis of E-cadherin and MUC2 expression in colon sections showed minor increases in the *EcN* WT group compared to the *EcN glyA* KO group (Extended Data Figure 2e,f). Finally, histological analysis of colon sections showed improved crypt morphology in the *EcN* WT group, with no significant changes in the number of goblet cells compared to the *EcN glyA* KO group (Extended Data Figure 2g). These findings indicate that while microbial folinic acid is sufficient to enhance epithelial homeostasis and suppress inflammation in immune-competent wild-type mice or mildly inflamed mice, its anti-inflammatory efficacy is lost in the chronic, advanced inflammatory states characteristic of IL10KO model.

Given our *in vitro* findings, we next investigated whether microbial folinic acid production could exert a more specific therapeutic effect on colon tumorigenesis. We hypothesized that the structural and epithelial stabilizing properties of folinic acid would provide an anti-neoplastic benefit that persists even in a highly inflammatory environment due to its inhibition of the pro-carcinogenic CAPN2 protease. To test this, we utilized an azoxymethane (AOM)-induced CRC model in IL10KO mice^39^ to determine if folinic acid suppresses tumorigenesis independently of its lack of efficacy in resolving chronic inflammation. Briefly, mice were injected intraperitoneally with AOM every week for the first 6 weeks, with concurrent biweekly gavages of *EcN* WT or *EcN glyA* KO bacteria for 14 weeks (Figure 5a). Fecal water content analysis, indicative of intestinal blockage or dysfunction^40^, showed a significant increase in the *EcN glyA* KO group compared to *EcN* WT group starting at week 11 (Figure 5b). Morphologically, colons from the *EcN glyA* KO group exhibited pronounced mucosal thickening and the presence of tumor-like nodular formations, whereas the *EcN* WT group displayed a more uniform and attenuated phenotype (Figure 5c). Colons from the *EcN glyA* KO group also exhibited significant architectural distortion and reduced lengths, further indicating severe structural damage and tumor-associated pathology compared to the *EcN* WT group (Figure 5c,d). Histological analysis of colon sections indicated a significant increase in the number of colon tumors and lymphovascular invasion events in the *EcN glyA* KO group compared to the *EcN* WT group (Figure 5e,f). Further, the *EcN glyA* KO group also displayed high scoring for tumor grade and mucosal invasion compared to the *EcN* WT group (Figure 5g,h). Fecal LCN-2 analysis confirmed that chronic inflammation remained unresolved across both groups (Extended Data Figure 3). An analysis of colonic CAPN2 protein levels showed a significant increase in the *EcN glyA* KO group compared to the *EcN* WT group indicative of increased CRC tumor burden (Figure 5i). Analysis of E-cadherin and MUC2 in colon sections showed a significant increase in the *EcN* WT group compared to the *EcN glyA* KO group, suggesting that in the context of CRC, folinic acid can improve gut epithelial integrity and barrier function (Figure 5j,k). Overall, these results suggest that while microbial folinic acid is insufficient to resolve chronic inflammation, it is sufficient to significantly attenuate CRC progression *in vivo* by inducing gut epithelial homeostasis.

**Fig. 5.**
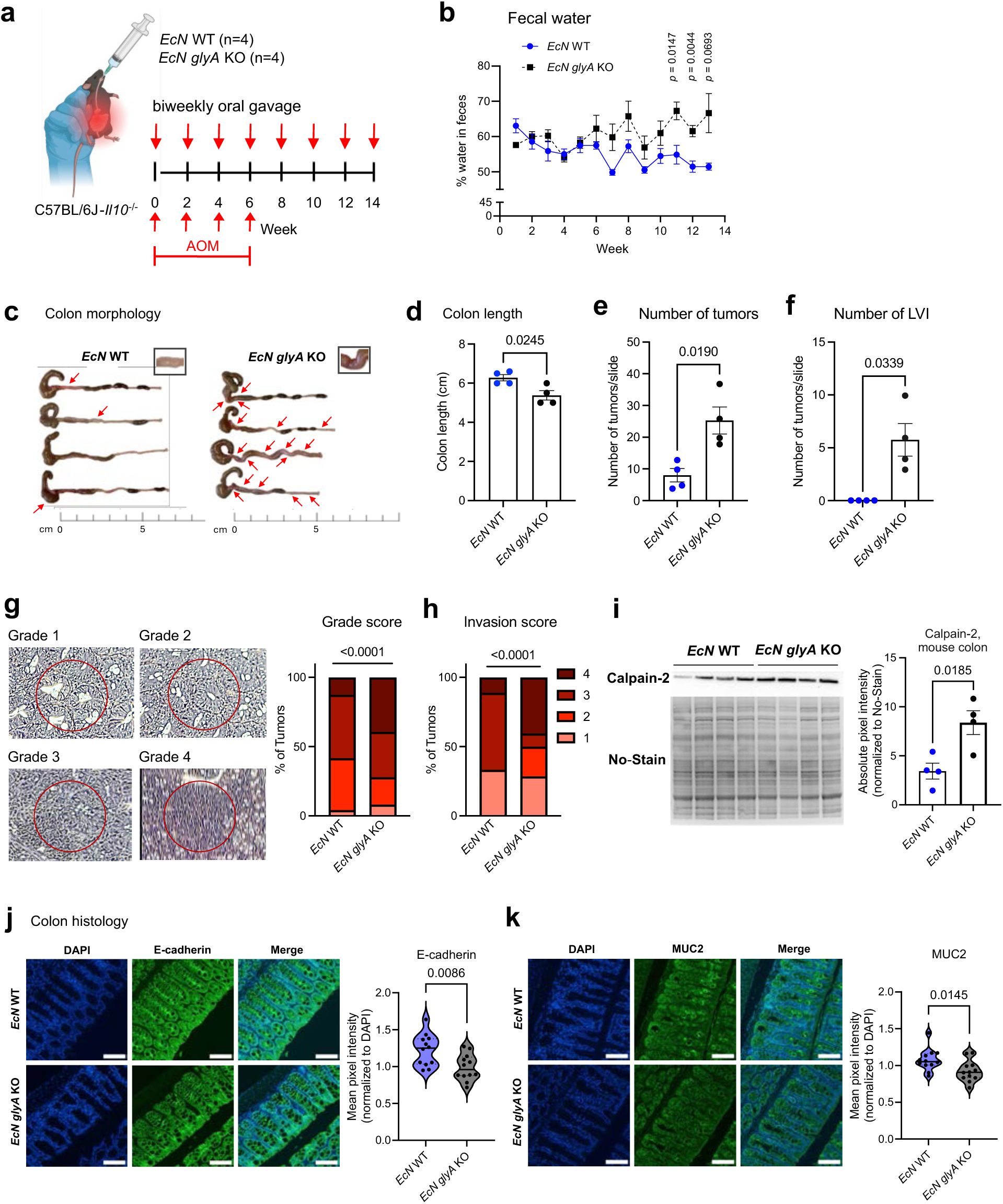
Microbial synthesis of folinic acid reduces CRC burden without mitigating inflammation. **a.** Schematic of weekly AOM treatment (10 mg/kg body weight) for six weeks and biweekly oral gavage of *E. coli* Nissle wild-type (WT) or *glyA* KO (10^9^ CFUs) administered to IL10KO mice at 16 weeks of age, followed by colon collection after 14 weeks. **b.** Fecal water content (n=4 per group, two-tailed Welch’s t test for multiple hypothesis testing). **c.** Colon images with tumor-like nodules and irregularities marked with red arrows. **d.** Colon length (cm) of each mouse. (n=4 per group, two-tailed Welch’s t test). **e, f**. Number of tumors (**e**) and lymphovascular invasion events (**f**) observed in histology slides. **g, h.** Tumor grade scoring system and scores (**g**) and tumor invasion scores (**h**). All images were scored in a blinded manner. **i.** Western blot of CAPN2 expression from mouse colons normalized to No-Stain. Each n represents one mouse (n=4 per group, two-tailed Welch’s t test). **j.** Representative images of mouse colons stained with E-cadherin and DAPI. Scale bar, 50 µm. Mean pixel intensity of E-cadherin was measured from 3 images per mouse colon using ImageJ and normalized to DAPI. (n=12 per group, two-tailed Welch’s test). **k.** Representative images of mouse colons stained with mucin 2 (MUC2) and DAPI. Scale bar, 50 µm. Mean pixel intensity of MUC2 was measured from 3 images per mouse colon using ImageJ and normalized to DAPI. (n=12 per group, Mann-Whitney test). All statistically significant *p* values are indicated in each graph. Data not marked are not statistically significant (*p* > 0.05). All bar graphs are represented as mean ± SEM, data points represent biological replicates.

### Microbial folinic acid suppresses CRC progression without major restructuring of the gut microbiota

To determine if microbial folinic acid production shapes overall microbiome community structure, we performed 16S rRNA sequencing across all *in vivo* experimental models (Figure 6). In WT mice, PCoA analysis revealed that colonization status significantly altered β-diversity with a minor shift in α diversity indicating that microbial folinic acid production is a key determinant of community assembly in homeostatic environment (Figure 6a-c). Phyla and Class level taxonomic abundances, and LEfSe analysis did not reflect distinct health- or disease-associated enrichments in either group, suggesting that folinic acid independently exerts beneficial effects on gut homeostasis and barrier integrity (Figure 6d-f). Interestingly, *EcN glyA* KO-colonized IL10KO mice exhibited significantly increased α-diversity compared to *EcN* WT controls, likely reflecting expansion of opportunistic taxa (Figure 6g-i). Phyla and Class level taxonomic abundances and LEfSe analysis confirmed that colonization with the folinic acid-deficient *EcN glyA* KO strain drives the microbiome toward an altered state, characterized by specific taxonomic shifts that are absent in the *EcN* WT-colonized or healthy models (Figure 6j-l). Strikingly, in the AOM-induced CRC model, we observed no significant differences in either α- or β-diversity between *EcN* WT and *EcN glyA* KO groups, further reflected in the phyla, class level abundances, and LEfSe analysis (Figure 6m-r). This suggests that in the context of chemically induced tumorigenesis, the protective anti-neoplastic effects of folinic acid are intrinsic to the metabolite’s signaling activity rather than a secondary consequence of community-wide microbiota remodeling.

**Fig. 6.**
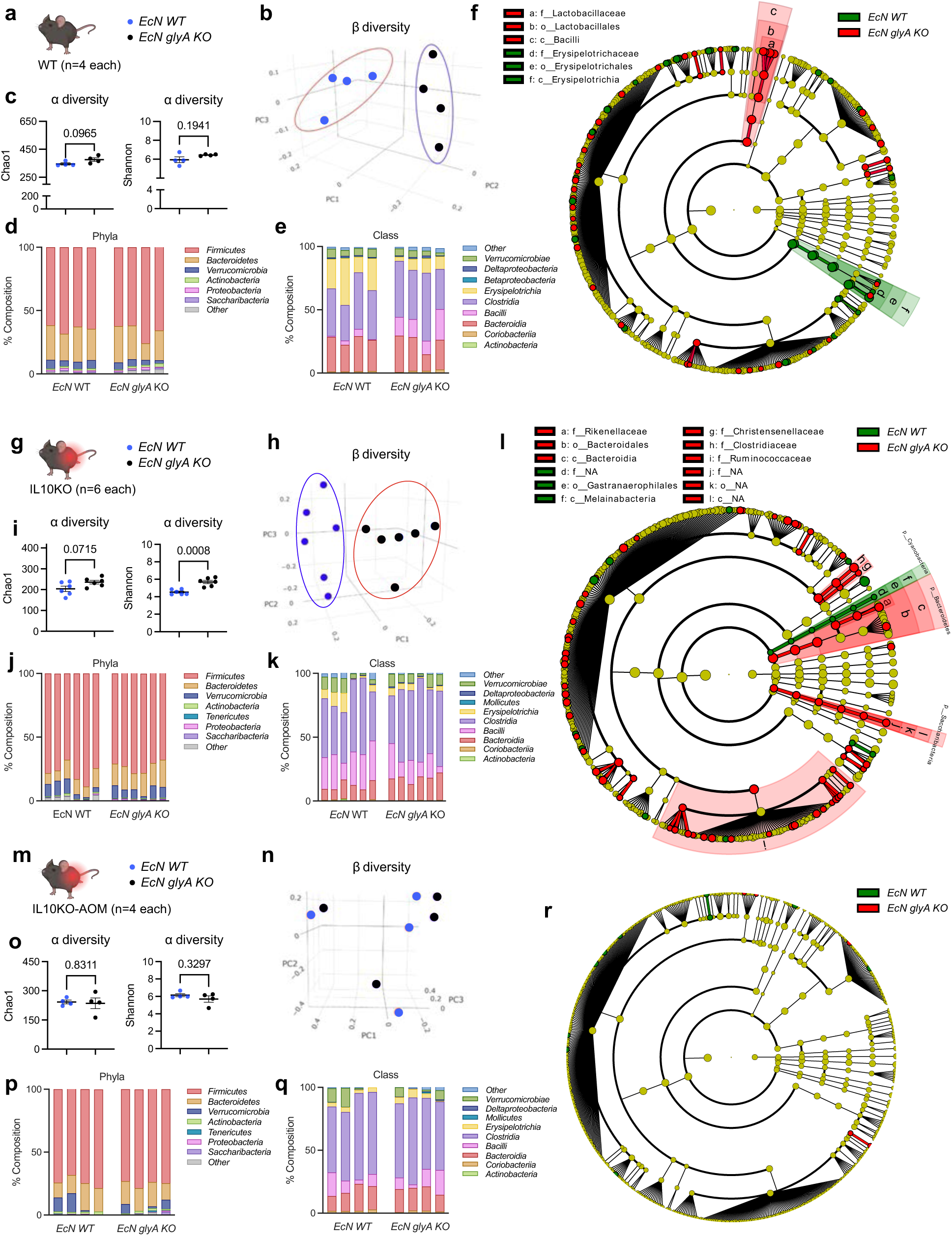
Microbial folinic acid suppresses CRC progression without major restructuring of the gut microbiota. **a.** *EcN* WT and *EcN glyA* KO gavaged WT mouse cecum samples analyzed by 16s rRNA sequencing (n=4 in each group). **b.** PCoA plot of Bray-Curtis β diversity. **c.** Chao and Shannon index α diversity (two-tailed Welch’s t test). **d, e.** Phyla (**d**) and Class (**e**) of bacterial taxa. **f.** LEfSe (Linear discriminant analysis Effect Size) cladogram analysis of microbiota. **g.** *EcN* WT and *EcN glyA* KO gavaged IL10KO mouse cecum samples analyzed by 16s rRNA sequencing (n=6 in each group). **h.** PCoA plot of Bray-Curtis β diversity. **i.** Chao and Shannon index α diversity (two-tailed Welch’s t test). **j, k.** Phyla (**j**) and Class (**k**) of bacterial taxa. **l.** LEfSe (Linear discriminant analysis Effect Size) cladogram analysis of microbiota. **m.** *EcN* WT and *EcN glyA KO* gavaged IL10KO + AOM mouse cecum samples analyzed by 16s rRNA sequencing (n=4 in each group). **n.** PCoA plot of Bray-Curtis β diversity. **o.** Chao and Shannon index α diversity (two-tailed Welch’s t test). **p, q.** Phyla (**p**) and Class (**q**) of bacterial taxa. **r.** LEfSe (Linear discriminant analysis Effect Size) cladogram analysis of microbiota. All statistically significant *p* values are indicated in each bar graph. Data not marked are not statistically significant (*p* > 0.05). All bar graphs are represented as mean ± SEM, data points represent biological replicates.

## Discussion

This study identifies a previously uncharacterized role for microbial folinic acid as an endogenous metabolite that actively suppresses colorectal cancer (CRC) progression. We show that folinic-acid producing bacteria are depleted in mouse models of colitis-associated CRC and in human clinical metagenomic cohorts with advancing disease severity. Our *in vitro* assays using HT29 cells show that treatment with folinic acid decreases metastasis, which is mediated through calpain-2 inhibition. Administration of a single microbe, wild-type *EcN*, which can produce folinic acid, improved epithelial and mucosal homeostasis and tumorigenesis in the colon compared to the *EcN glyA* KO strain. While *EcN* has been previously documented to improve CRC outcomes, our findings demonstrate that its protective efficacy is dramatically attenuated when its ability to produce folinic acid is genetically abolished via *glyA* deletion.

The clinical utility of folinic acid is well-established; owing to its chemical stability and pharmacokinetics, it is the folate of choice for adjuvant chemotherapy, where it stabilizes the 5-fluorouracil (5-FU) bond with thymidylate synthase^41,42^. However, this heightened potency is often accompanied by severe gastrointestinal side effects, including inflammation, damage, drug resistance, and gut dysbiosis^43^. Because these outcomes typically occur in patients receiving highly cytotoxic co-therapies (e.g., fluoropyrimidine, methotrexate), it has historically been difficult to discern whether folinic acid is causative or correlative towards these outcomes. There have been no reports on the effects of folate production from the microbiome in CRC therapeutics and outcomes. Our data suggests a paradigm shift, wherein increasing local colonic concentrations of folinic acid via targeted microbial production may provide therapeutic benefits while circumventing the systemic toxicity associated with high-dose pharmacological supplementation. Our findings also highlight the role of calpain-2 (CAPN2) in this pathway. We demonstrate that folinic acid suppresses CRC metastasis by inhibiting CAPN2, an intracellular cysteine protease known to promote tumor progression and therapy resistance in numerous cancers including lung, breast, pancreatic, and colorectal cancer^44–46^. While targeted pharmacological inhibition of CAPN2 has been shown to attenuate tumor growth and reduce metastasis^47^, the ubiquitous expression of CAPN2 in mammalian tissues often leads to adverse, untargeted effects. By leveraging the gut microbiome as a localized bioreactor, we may achieve the anti-metastatic effects of CAPN2 inhibition with greater precision.

Folate cannot be synthesized naturally by humans and must be acquired through diet, supplementation, or microbial production^48^. Our study underscores the importance of the microbiome in this metabolic supply chain, as folate-deficient diets may exacerbate CRC progression by limiting microbial folinic acid reservoirs^49^. Moving forward, characterizing the precise bacterial consortia, such as *Lactiplantibacillus*, *Lactococcus*, *Bifidobacterium*, and *Streptococcus* capable of high-level folinic acid production, will be essential for developing next-generation probiotic therapeutics. Future research should also investigate whether patient-specific microbiome profiles, specifically those enriched for folate-producing strains, correlate with improved chemotherapy response and long-term CRC remission. This study potentially offers a new approach into colorectal cancer therapy by altering the microbiome to produce beneficial metabolites, such as folinic acid, to maintain epithelial and mucosal integrity within the host.

## Methods

### Mouse studies

C57BL/6J male mice (6 weeks old) and C57BL/6J-*Il10^-/-^* (IL10KO) male and female mice (11-16 weeks old; B6.129P2-Il10tm1Cgn/J) were purchased from Jackson Laboratory (Bar Harbor, ME) and bred in the University of Wisconsin-Madison Biomedical Research Model Services (BRMS) facility. Animals are transferred to the Chaudhari lab animal facility for experiments. Germ-free male mice (6-8 weeks old) were obtained from the National Gnotobiotic Rodent Resource Center (NGRRC) at the University of North Carolina at Chapel Hill and bred in the University of Wisconsin-Madison BRMS Gnotobiotic Shared Resource (GSR) facility. Germ-free animals were maintained in a controlled environment in plastic flexible film gnotobiotic isolators.

Each mouse was orally gavaged with 100 µL of either *EcN* WT or *glyA* KO at 10^10^ CFU/mL diluted in 1x DPBS. This treatment was repeated biweekly for the entirety of each experiment.

All animals were housed in climate-controlled animal facilities under a 12 h light/12 h dark cycle with ad libitum access to food and water. Mice were fasted for 4 h prior to sacrifice. Tissues and fecal samples were immediately snap-frozen on dry ice and stored at -80°C until further analysis. For serum, blood was allowed to clot at room temperature for 30 min and centrifuged at 1000 x *g* for 20 min. The supernatant was transferred to microcentrifuge tubes and stored at −80°C until further analysis. All procedures were approved by the Institutional Animal Care and Use Committee (IACUC) at the University of Wisconsin-Madison.

### Folate metabolomics

#### Bacterial folate extraction

*Escherichia coli* Nissle 1917 (*EcN*), was streaked onto Luria broth (LB) plates (10 g/L tryptone, 5 g/L yeast extract, 5 g/L NaCl, 15 g/L agar) from 25% glycerol stocks and incubated aerobically at 37°C for 16–18 h. For *EcN glyA* KO, LB plates with kanamycin (25 µg/mL) were used. Single colonies were used to inoculate 10 mL culture tubes containing 5 mL of liquid LB. Cells were grown overnight at 37°C at 150 revolutions per minute (RPM), then subcultured into dilute LB media (1:9 LB:M9 media) to mid-log phase (M9 6 g/L Na2HPO4, 3 g/L KH2PO4, 5 g/L NaCl, and 1 mM MgSO4).

After reaching mid-log phase, cultures were pelleted via centrifugation at 4000 x *g* for 5 minutes, media was removed and pellets weighed. After weighing, 200 μL of cold bacterial extraction buffer (20 mM ascorbic acid, 20 mM BME, 50:50 methanol and water) was used to resuspend each pellet and transferred to a 2-mL bead-homogenizing tube (Fisher Scientific, 02681374) with glass beads (Bertin Technologies, P000928-LYSK0-A.0). Samples were homogenized for 1 min at 4 m/s (Bead Ruptor Elite, Omni International) before being centrifuged at 18200 x *g* for 10 min at 4°C. After centrifugation, the supernatant of each sample was transferred to a clean 1.5 mL Eppendorf tube. Samples were dried down in a SpeedVac (Thermo Scientific, Savant SPD120 and Savant UVS450) at 35°C for 90 min.

After drying, samples were resuspended in 200 μL folate extraction buffer (20 mM ascorbic acid, 20 mM ammonium acetate and 20 mM BME, pH 8.1, adjusted with NaOH, prepared fresh) with spiked internal standards. Extracts were kept on ice unless noted otherwise. To each sample, 5 μL of pooled rat serum (Santa Cruz Biotechnology, sc-45135) was added and sample was vortexed briefly to mix, before incubating at 37°C for 45 min. After incubation, tubes were stored on ice before being boiled at 100°C for 5 min. Samples were centrifuged at 18200 x *g* for 5 min to pellet the remaining debris.

Supernatants were filtered by solid-phase extraction (SPE) using a Strata-X 33-μm polymeric reverse-phase 96-well SPE plate (Phenomenex, 8E-S100-AGB). SPE wells were conditioned using 200 μL of methanol and equilibrated using 200 μL of resuspension buffer (20 mM ammonium acetate and 20 mM BME) by centrifugation at 3000 x *g* for 30 seconds after each solvent addition. Sample supernatants were passed through the column, and wells were rinsed using 200 μL of resuspension buffer. SPE plates were moved to a clean 96-well plate, and metabolites were eluted using two passthroughs of 100 μL elution buffer (50:50 methanol and water, 20 mM ammonium acetate and 20 mM BME). Eluted samples were dried down in a SpeedVac (Thermo Scientific, Savant SPD120 and Savant UVS450) at 35°C for 90 min. Dried down samples were resuspended in 40 μL of resuspension buffer and were moved to autosampler vials (Fisher Scientific, 3452232) for LC–MS analysis. Ascorbic acid and ammonium acetate for folate extraction were purchased from Santa Cruz Biotechnology (Santa Cruz Biotechnology, sc-202686 and sc-203818). BME was purchased from MP Biomedicals (MP Biomedicals, 190242). Water for folate extraction was obtained from a GenPure Pro water purification system (Thermo Scientific, 50131948).

#### Fecal and cecum folate extraction

Frozen cecum tissue was cut on dry ice or one frozen fecal pellet per mouse were weighed for extraction. Approximately 20-50 mg of tissue was used for extraction. Sample was added to a 2-mL bead-homogenizing tube (Fisher Scientific, 02681374) with glass beads (Bertin Technologies, P000928-LYSK0-A.0). Then, 200 μL (for samples <50 mg) or 400 μL (for samples >50 mg) of folate extraction buffer (20 mM ascorbic acid, 20 mM ammonium acetate and 20 mM β-mercaptoethanol (BME), pH 8.1, adjusted with sodium hydroxide, prepared fresh) with spiked internal standards was added to each tissue sample and homogenized for 1 min at 4 m/s (Bead Ruptor Elite, Omni International). Homogenates were kept on ice unless noted otherwise. Homogenates were centrifuged at 18200 x *g* for 5 min at 4°C. Pooled rat serum (Santa Cruz Biotechnology, sc-45135), 5 μL (for samples <50 mg) or 10 μL (for samples >50 mg), was added to each sample, followed by briefly vortexing to mix and incubating at 37°C for 45 min. After incubation, tubes were stored on ice before being boiled at 100°C for 5 min. Samples were centrifuged at 18200 x *g* for 5 min to pellet the remaining debris. Supernatants were filtered by SPE using a Strata-X 33-μm polymeric reverse-phase 96-well SPE plate (Phenomenex, 8E-S100-AGB). After each solvent addition, samples were passed through the column by centrifugation at 3000 x *g* for 30 s. SPE wells were conditioned using 200 μL of methanol and equilibrated using 200 μL of resuspension buffer (20 mM ammonium acetate and 20 mM BME, pH 8.1). Sample supernatant was passed through the column and wells were rinsed using 200 μL of resuspension buffer. SPE plates were moved to a clean 96-well plate, and metabolites were eluted using two passthroughs of 100 μL elution buffer (50:50 methanol and water, 20 mM ammonium acetate and 20 mM BME). Eluted samples were dried down in a SpeedVac (Thermo Scientific, Savant SPD120 and Savant UVS450) at 35°C for 90 min. Dried down samples were resuspended in 40 μL of resuspension buffer and were moved to autosampler vials (Fisher Scientific, 3452232) for LC–MS analysis. Ascorbic acid and ammonium acetate for folate extraction were purchased from Santa Cruz Biotechnology (Santa Cruz Biotechnology, sc-202686 and sc-203818). BME was purchased from MP Biomedicals (MP Biomedicals, 190242). Water for folate extraction was obtained from a GenPure Pro water purification system (Thermo Scientific, 50131948).

#### LC-MS

LC–MS metabolomics was performed using a Vanquish ultrahigh-performance LC system (Thermo Scientific) coupled to an Orbitrap Exploris 120 MS instrument (Thermo Scientific). Chromatography was performed using a reverse-phase polar C18 column (Phenomenex, 00D-4748-AN; Luna Omega Polar C18, 1.6-µm particle size, 2.1 × 100 mm column). The column chamber and preheater were set to 40°C. Solvent A was 95% H2O and 5% methanol with 5 mM DMHA (Acros Organics, 314740250) pH 8.1, adjusted with formic acid. Solvent B was 100% methanol with 5 mM DMHA. All solvents were LC–MS grade purchased from Fisher Chemical (Thermo Scientific, CAS 67-56-1). The total run time was 12 min. Flow rate was held constant at 0.4 mL/min. The chromatography gradient was as follows: 5% solvent B for 0.5 min, linear increase to 85% B over 5.2 min, increase to 100% B for 1.3 min and decrease to 5% to equilibrate the column for 4 min. Eluent from the column was analyzed by MS from the start of the run until 5 min, after which flow was directed to waste for the remainder of the run. Ionization was performed using an HESI source in negative mode at 2,500 V. Nitrogen gas flow was set to 50 (sheath), 10 (auxiliary) and 1 (sweep) in arbitrary units. The ion transfer tube was maintained at 325°C with the vaporizer at 350°C. Quantification was performed using Freestyle version 1.8.65.0 (Thermo Fisher) and MAVEN78 version 2.10.21 to identify and calculate integrated peak areas. Manual curation confirmed peak identification and parameters. All signals were normalized to internal standards (methotrexate, folic acid-d2) for injection variation. Identification of folates without available standards was performed using SIRIUS version 6.3.3 and comparisons to previously published results.

#### *glyA* counts

Metagenomic data were downloaded from NCBI. Quality of sample sequences from each dataset was checked with FastQC. None of the chosen datasets showed significant quality control issues. Metagenomes were assembled using SPAdes in metagenome mode. Then bowtie2 was used to align reads to the assembled metagenomes. Bowtie-build was run to build a reference for the metagenome, followed by bowtie2. Samtools was used to convert the resulting bam file to a sam file. Picard tools were then used to identify duplicate reads. Prokka was used to annotate the genomes. Prokka was run with the metagenome and compliant options. A bash script was then used to process prokka output to remove lines containing “#” and then retain lines only with “ID=”. Any metagenomes that resulted in errors when running any of these software were discarded from future analysis.

A custom script, htseq_count.py, was used to apply HTSeq to determine the number of counts assigned to each gene. A second custom script, compare_gene_abudnaces.py, was used to determine the counts assigned to the gene of interest divided by the total reads mapped to a gene. This normalized count was then used to compare abundance of the gene of interest across metagenomic samples with disease state determined by metadata from the original study.

#### SHMT sequence slignment

To screen SHMT protein sequence similarity across 119 gut bacterial species (hCom2)^24^, SHMT from *E. coli* (UniProt accession: <u>NP_417046.1</u>) was used as a query for protein-protein BLAST (BLASTP) searches using the NCBI BLAST against the RefSeq Select proteins database. Word size was set to 6 and default scoring parameters were applied. For each bacterial species, the best hit based on the lowest E-value (expectation value) was selected, and the corresponding maximum alignment score was used for comparative analysis and visualization in a heatmap.

#### 16S rRNA sequencing

16S rRNA gene sequencing was performed by Zymo Research (Irvine, CA) using the ZymoBIOMICS® Targeted Sequencing Service. Genomic DNA was extracted using the ZymoBIOMICS®-96 MagBead DNA Kit or ZymoBIOMICS® DNA Miniprep Kit according to the manufacturer’s protocols. The V3–V4 region of the bacterial 16S rRNA gene was amplified and libraries were prepared using the Quick-16S™ NGS Library Prep Kit (Zymo Research). Libraries were sequenced on an Illumina NextSeq platform (600-cycle kit). Amplicon sequence variants (ASVs) were inferred using the DADA2 pipeline (Callahan et al.^50^), and chimeric sequences were removed during processing. Taxonomic assignment and diversity analyses were performed using QIIME v1.9.1 (Caporaso et al.^51^) with the Zymo Research reference database.

#### In vitro assays

HT29 (ATCC HTB-38, USA) cells were seeded at a density of 2.0 × 10^6^ cells per 75-cm2 flask. The cells were grown in tissue culture flasks at 37°C, 5% CO_2_. The cells were sub-cultured at ∼ 90% confluence (∼6 days) by 0.25% trypsin and 0.02% EDTA solution. DMEM (GenClone, 25-500) was supplemented with 10% fetal bovine serum and 1% penicillin and streptomycin (complete media).

#### CFU assay

Seeded HT29 cells at 100 cells per well in a 6-well plate and treated with 25 µM folinic acid in DMSO or 0.5% DMSO and incubated at 37°C, 5% CO_2_ for 7 days, exchanging the media and treatment every 3-4 days. Gently washed the cells with 1 mL of 1x DPBS, added 1 mL of 0.1% crystal violet in 20% methanol per well and incubated at room temperature for 10 min, then gently washed 3 x with 1x DPBS. CFUs were imaged using an iBright imaging system (Invitrogen). The volume/area of each colony was calculated from the Colony Count Analysis Data Table generated by the iBright instrument using the Universal, Visible Crystal Violet settings.

#### Immunocytochemistry

HT29 cells were seeded at 40,000 cells/well in 100 µL on 96-well glass-bottom plates (Cellvis, NC0536760) and treated with 25 µM folinic acid in DMSO or 0.5% DMSO for 10 days, exchanging the media and treatment every 3-4 days. Cells were washed three times with 1x PBS and blocked in PBS containing 4% donkey serum (GeneTex, GTX27475) and 0.3% Triton X-100 for 1 h at room temperature with gentle agitation. Cells were incubated with primary antibodies diluted in blocking solution overnight at 4°C with gentle agitation. The following day, cells were washed three times in PBS (5 min per wash) at room temperature and incubated with fluorophore-conjugated secondary antibodies diluted in blocking solution for 1 h at room temperature, protected from light. After washing with PBS, nuclei were stained with DAPI (1:3000 in PBS) for 5 min. Cells were washed again with PBS before imaging. Images were aquired using appropriate filter sets on a Nikon A1R+ confocal microscope at the Biochemistry Optical Core (BOC) Facility at UW–Madison. Primary antibody used for immunocytochemistry fluorescence staining was E-cadherin (1:1000; Cell Signaling Technology). Goat anti-rabbit IgG (H+L) Cross-Adsorbed Alexa Fluor™ 488 (1:500, Thermo Fisher) was used as a secondary antibody.

#### Western blot analysis

Mouse colon tissue (∼30 mg) or HT29 cells were lysed in 100-200 µL of ice-cold 1x Cell Lysis Buffer (Cell Signaling Technology) supplemented with protease inhibitor cocktail (Roche). Tissue was homogenized using a Bead Ruptor Elite (Omni International, Kennesaw, GA). Samples were centrifuged at 18200 x *g* for 10 min at 4°C, and supernatants were collected. Protein concentrations were determined using a Bradford assay (Thermo Scientific) with bovine serum albumin (BSA) as a standard. Proteins were mixed with 4x loading buffer supplemented with β-mercaptoethanol, boiled at 100°C for 10 min, and separated on precast Bis-Tris SDS-PAGE gels (Invitrogen). Proteins were transferred onto PVDF membranes using wet transfer at 30 V for 1 h. Membranes were blocked in StartingBlock™ Blocking Buffer (Thermo Scientific) for 1 h at room temperature and incubated with primary antibodies diluted in 5% BSA in TBS-T overnight at 4°C. After washing with TBS-T, membranes were incubated with HRP-conjugated secondary antibodies diluted in 5% BSA in TBS-T for 1 h at room temperature. Blots were developed using SuperSignal™ West Pico PLUS Chemiluminescent Substrate (Thermo Scientific). Blot images were obtained from iBright imaging system (Invitrogen). Band intensities were quantified using the iBright imaging system (Invitrogen) with automated band detection and normalized to total protein. Primary antibodies used for western blot are: E-cadherin (1:1000; Cell Signaling Technology) and Calpain-2 (1:1000; Cell Signaling Technology). Mouse anti-rabbit IgG-HRP (1:5000; Santa Cruz) was used as a secondary antibody.

#### Transwell assays

HT29 cells were seeded onto 8.0 µm pore-size inserts in 24-well plates (Costar 3422) at a density of 50,000 cells in 200 µL of serum-free media on the apical side and 500 µL of complete media on the basolateral side. Cells were immediately treated with 25 µM folinic acid (FA) in DMSO or 0.5% DMSO. Basolateral and apical media with FA or DMSO were exchanged every 3-4 days. Following treatment, transwell inserts were removed and transferred to a new 24-well plate. Cells were gently washed x 1 with 1x DPBS and stained with 0.1% crystal violet in 20% methanol on the basolateral side, as previously described. Apical cells attached to the transwell insert were fixed with 10% neutral buffered formalin, washed 3 x with 1x PBS, and stained with E-cadherin and DAPI as previously described.

To generate enough mesenchymal cells to plate onto new transwell plates, HT29 cells were seeded onto 8 µm pore-size inserts in 6-well plates (Costar 3428) at a density of 200,000 cells per insert in 1 mL of serum-free media and incubated at 37°C, 5% CO_2_ for ∼12 days. When enough cells had migrated and formed colonies on the basolateral side, the mesenchymal cells were transferred to new transwell plates (Costar 3422) and treated with 25 µM folinic acid (FA) in DMSO or 0.5% DMSO, as previously described.

#### Electron microscopy

HT29 cells in transwells were fixed in a modified Karnovsky’s fixative (2.5% glutaraldehyde/2.0% formaldehyde in 0.1 M sodium phosphate buffer (PB), pH=7.2), post fixed in 1.0% osmium tetroxide in 0.1M PB, dehydrated in a grade series of ethanol, transitioned in acetone and embedded in Embed 812 (EMS, Morgantown, PA). Polymerized samples were sectioned (80nm) on an ultramicrotome and sections were placed onto formvar coated 2x1 Cu slot grids and post-stained with uranyl acetate and lead citrate. The sections were viewed at 80kV on a FEI CM120 TEM and imaged on an AMT (Danvers, MA) BioSprint12 digital camera.

### Target engagement proteomics

#### Lysate-based proteome integral solubility alteration

Frozen HT29 cell pellets were thawed on ice and resuspended in lysis buffer (1 X PBS pH 7.4, 1 mM MgCl_2_, protease inhibitor). The proteomes were extracted using a dounce homogenizer (20 strokes). The extracts were spun at 300 x *g* for 3 min to remove any unbroken cells. The resulting crude extract was diluted to 2 mg/mL in lysis buffer. Each compound was added to lysis buffer at a 2 X concentration. In order to initiate the experiment, an equal volume of crude extract and treatment buffer were combined, to achieve a final protein concentration of 1 mg/mL and compound concentration of 10 μM, and incubated for 30 min. After incubation, an equal volume of each sample was transferred to 10 PCR tubes. The PCR tubes were heated across a thermal gradient ranging from 48°C to 58°C for 3 min to induce thermal denaturation. An equal volume from each PCR tube was pooled. An equal volume of extraction buffer (1 X PBS pH 7.4, 1% NP-40, protease inhibitors) was added to added to each pooled sample to achieve a final NP-40 concentration of 0.5%. Samples were incubated for 10 min at 4°C on a roller. Extracted samples were spun at 21,000 x *g* for 90 min to separate insoluble aggregates from soluble protein. An equal volume from each soluble fraction was collected and prepared for LC-MS/MS analysis.

#### LC-MS/MS sample preparation

Samples (15-20 μg protein) were diluted in prep buffer (400 mM EPPS pH 8.5, 1% SDS, 10 mM tris(2-carboxyethyl)phosphine hydrochloride) and incubated at room temperature for 10 min. Iodoacetimide was added to a final concentration of 10 mM to each sample and incubated for 25 min. in the dark. Finally, DTT was added to each sample to a final concentration of 10 mM. A buffer exchange was carried out using a modified SP3 protocol^52^. Briefly, ∼250 μg of each SpeedBead Magnetic Carboxylate modified particles (Cytiva; 45152105050250, 65152105050250) mixed at a 1:1 ratio were added to each sample. 100% ethanol was added to each sample to achieve a final ethanol concentration of at least 50%. Samples were incubated with gentle shaking for 15 min. Samples were washed three times with 80% ethanol. Protein was eluted from SP3 beads using 200 mM EPPS pH 8.5 containing trypsin (ThermoFisher Scientific) and Lys-C (Wako). Samples were digested overnight at 37°C with vigorous shaking. Acetonitrile was added to each sample to achieve a final concentration of 30%. Each sample was labelled, in the presence of SP3 beads, with ∼65 μg of TMTpro-16plex reagents^53,54^ (ThermoFisher Scientific). Experimental layouts for each experiment were described in corresponding source data tables. Following confirmation of satisfactory labelling (>97%), excess TMTpro reagents were quenched by addition of hydroxylamine to a final concentration of 0.3%. The full volume from each sample was pooled and acetonitrile was removed by vacuum centrifugation for one hour. The pooled sample was acidified using formic acid and peptides were de-salted using a Sep-Pak Vac 50 mg tC18 cartridge (Waters). Peptides were eluted in 70% acetonitrile, 1% formic acid and dried by vacuum centrifugation. The peptides were resuspended in 10 mM ammonium bicarbonate pH 8, 5% acetonitrile and fractionated by basic pH reverse phase HPLC. In total 24 fractions were collected. The fractions were dried in a vacuum centrifuge, resuspended in 5% acetonitrile, 1% formic acid and desalted by stage-tip. Final peptides were eluted in, 70% acetonitrile, 1% formic acid, dried, and finally resuspended in 5% acetonitrile, 5% formic acid. In the end, 12 of 24 fractions were analyzed by LC-MS/MS.

#### Mass spectrometry data acquisition

Data were collected on an Orbitrap Eclipse mass spectrometer (ThermoFisher Scientific) coupled to a Proxeon EASY-nLC 1000 LC pump (ThermoFisher Scientific). Peptides were separated using a 120-min gradient at 500 nL/min on a 30-cm column (i.d. 100 μm, Accucore, 2.6 μm, 150 Å) packed in house. High-field asymmetric-waveform ion mobility spectroscopy (FAIMS) was enabled during data acquisition with compensation voltages (CVs) set as −40 V, −60 V, and −80 V. MS1 data were collected using the Orbitrap (60,000 resolution; maximum injection time 50 ms; AGC 4 × 10^5^). Determined charge states between 2 and 6 were required for sequencing, and a 60 s dynamic exclusion window was used. Data dependent mode was set as cycle time (1 s). MS2 scans were performed in the Orbitrap with HCD fragmentation (isolation window 0.5 Da; 50,000 resolution; NCE 36%; maximum injection time 86 ms; AGC 1 × 10^5^).

#### Mass spectrometry data analysis

Raw files were first converted to mzXML, and monoisotopic peaks were re-assigned using Monocle. Database searching included all human entries from Uniprot (downloaded in February 2020). The database was concatenated with one composed of all protein sequences in the reversed order. Sequences of common contaminant proteins (e.g., trypsin, keratins, etc.) were appended as well. Searches were performed using the comet search algorithm. Searches were performed using a 50-ppm precursor ion tolerance and 0.02 Da product ion tolerance. TMTpro on lysine residues and peptide N termini (+304.2071 Da) and carbamidomethylation of cysteine residues (+57.0215 Da) were set as static modifications, while oxidation of methionine residues (+15.9949 Da) was set as a variable modification.

Peptide-spectrum matches (PSMs) were adjusted to a 1% false discovery rate (FDR). PSM filtering was performed using linear discriminant analysis (LDA) as described previously^55^ while considering the following parameters: comet log expect, different sequence delta comet log expect (percent difference between the first hit and the next hit with a different peptide sequence), missed cleavages, peptide length, charge state, precursor mass accuracy, and fraction of ions matched. Each run was filtered separately. Protein-level FDR was subsequently estimated at a data set level. For each protein across all samples, the posterior probabilities reported by the LDA model for each peptide were multiplied to give a protein-level probability estimate. Using the Picked FDR method^56^, proteins were filtered to the target 1% FDR level.

For reporter ion quantification, a 0.003 Da window around the theoretical *m/z* of each reporter ion was scanned, and the most intense *m/z* was used. Reporter ion intensities were adjusted to correct for the isotopic impurities of the different TMTpro reagents according to manufacturer specifications. Peptides were filtered to include only those with a summed signal-to-noise (SN) of 160 or greater across all channels. For each protein, the filtered peptide TMTpro SN values were summed to generate protein quantification.

#### CAPN2 protein expression and purification

CAPN2 catalytic subunit expressing pET-28a plasmid was generated by Craig Bingman at the University of Wisconsin-Madison Biochemistry Crystal Core and transformed into KRK cells (Promega, Madison WI). A single colony of KRK cells expressing recombinant CAPN2 with N-terminal 6xHis tag in pET-28a plasmid was inoculated in 5 mL LB with kanamycin and grown overnight at 37°C with shaking. The following day, 2 mL culture was expanded into 50 mL and protein expression was induced with 0.2 mM IPTG and 10 mM rhamnose. The culture was grown at 37°C overnight with shaking. The next day, 1 mL culture was removed to assess protein expression. Both the 1 mL aliquot and the remaining 49 mL culture were pelleted by centrifugation at 8000 x *g* for 15 min. Pellets from 49 mL were flash frozen and stored in -80°C pending confirmation of protein expression.

The 1 mL pellets were resuspended in 100 µL lysis buffer and sonicated in 3 x 15 s pulses, with 1 min intervals on ice between pulses. Lysates were clarified by centrifugation at 8000 x *g* for 5 min. 20 µL lysates were mixed with 6 µL Laemmli sample buffer containing NuPAGE sample reducing agent (Thermo Fisher), and boiled at 100°C for 5 min. The entire sample volume was subjected to western blot analysis for confirmation of protein expression. Blots were probed with mouse anti-His primary antibody (Proteintech, 66005-1-Ig) overnight at 4°C, followed by goat anti-mouse IgG HRP secondary antibody (Invitrogen, 62-6520) for 1 h at RT. Blot was developed with ECL reagent and imaged on the iBright CL1500 Imaging System.

Upon confirmation of protein expression from the 1 mL culture, the remaining 49 mL frozen pellet was thawed and lysed by sonication in 5 mL lysis buffer as described above. Cell lysate was clarified by centrifugation at 15,000 x *g* for 15 min. Clarified lysate was incubated with 1 mL cOmplete™ His-Tag Purification Resin (Roche) (50% slurry in IMAC Wash Buffer) for 1 h at 4°C on a tube roller. The mixture was drained through gravity-flow columns, and the resin was washed twice with 10 mL IMAC Wash Buffer and incubated with 1 mL IMAC Elution Buffer for 5 min before elution. The eluted protein was immediately mixed with 10 µL Mg:EDTA chelating solution to prevent self-cleavage. Protein expression was validated using No-Stain Labelling Reagent (Invitrogen, A44717) as described above.

Prior to the 5-f-THF binding assay and protein quantification, the eluate was buffer-exchanged into IMAC Equilibration Buffer using an Amicon® Ultra Centrifugal Filter (10 kDa MWCO, Millipore, UFC801008), with eight rounds of dilution/concentration in 5 mL IMAC Equilibration Buffer, to reduce the amount of Mg:EDTA and imidazole that could interfere with these steps.

#### CAPN2 binding assays

Purified CAPN2 catalytic subunit (100 mg/mL) was used in each replicate of the binding assay in 100 µL of binding buffer (25 mM HEPES, 100 mM NaCl, 0.01% Triton X-100, 5% glycerol, 1 mM DTT, 0.2% ascorbic acid, pH 7.25). CAPN2 was incubated with equimolar concentrations of folates (10 µM) at 37°C for 30 min. Protein was purified using IMAC purification as described above, and subjected to folate metabolomics analysis for quantification of folates. For CAPN2 displacement assay, purified protein was incubated with increasing concentrations of folinic acid at 37°C for 30 min in binding buffer. Bound protein was purified using IMAC purification, followed by incubation with 100 nM of CAPN2i at 37°C for 30 min in binding buffer. Displaced folinic acid was separated from protein using IMAC purification and measured using folate metabolomics LC-MS.

#### CAPN2KO cell line generation

A gene knockout kit (EditCo Bio, Redwood City, CA, USA) containing three chemically synthesized gRNAs (Spacer sequences: AGTTCTGGCAATACGGCGAG, GAGCAGCTCCCCGTCCTTGG, GGCGTATGCCTTCTCCAGCA) targeting the CAPN2 gene and a SpCas9 nuclease-recombinant protein (EditCo Bio, Redwood City, CA, USA) were used. The Neon Transfection System (Thermo Fisher, Waltham, MA, USA) was used to electroporate the gRNAs complexed with SpCas9 into the HT29 cells. After 72 h, genomic DNA was extracted using QuickExtract™ DNA Extraction Solution (LGC Genomics LLC, Alexandria, MN, USA). The CAPN2 gene was amplified using primers (IDT, Coralville, IA, USA). Editing of the CCR5 gene was measured using Sanger sequencing analyzed using the ICE Analysis software (EditCo Bio, Redwood City, CA, USA) to confirm gene knockout.

#### CAPN2OE cell line generation

CAPN2OE cell line was generated as described previously^29^. Briefly, HT29 cells were incubated with 1 nM of SN-38 (Sigma, H0165) for 24 h in complete DMEM media followed by recovery in drug-free buffer for ∼2 weeks. Dose of drug was doubled in every consecutive exposure, until resistant cells were obtained that could divide in 60 µM SN-38. Cells were allowed to recover in complete DMEM for 2 weeks prior to use in any experiment.

#### *E. coli* Nissle *glyA* deletion and *in vivo* quantification

The lambda red recombination system was used to delete the *glyA* gene from *EcN* using a method described previously^34^ with the following oligos: 5’ – GCTAAAGCTTTTAAGAAGGAGATATACATATGTAAAGCGTGAAATGAACATTGCCG – 3’ and 5’ – GCTATCTAGATTATGCGTAAACCGGGTAACGTGC – 3’. *In vivo* quantification of *EcN* was performed in fecal samples using *EcN* specific primers: (5’ – ATACTACGACGGTAAATGGT – 3’ and 5’ - TACATCAGTATCGGTAGCAT – 3’); and normalized to bacterial 16S primers (5’ – TATGGTAATTGTGTGCCAGCMGCCGCGGTAA – 3’; and 5′ – AGTCAGTCAGCCGGACTACHVGGGTWTCTAAT – 3’). Feces samples pre- and post-colonization were weighed out and diluted in 1x PBS to a concentration 5 mg/mL and 0.5 mg/mL. A qPCR was performed on these samples in triplicate using SYBR Green Master Mix (Thermo Fisher, Waltham, MA, USA) and run on the QuantStudio (Thermo Fisher, Waltham, MA, USA). Primers (IDT, Coralville, IA, USA) amplifying the *EcN* fimA gene were used to measure the amount of *EcN* in the feces. Primers amplifying the 16S gene were used to measure the amount of all bacteria in the feces.

#### H&E and immunofluorescent staining

Formalin-fixed and paraffin-embedded colon tissues were sectioned at 2-5 µm thickness and dried at 65°C for 25 min prior to staining with hematoxylin and eosin (H&E), E-cadherin, or mucin 2 (MUC2) using the Leica ST5010 Autostainer XL (Leica Biosystems) according to the UWCC Experimental Animal Pathology Laboratory protocol. H&E sections were sequentially treated with xylene, graded ethanol, Hematoxylin MX 560, Define MX, Blue Buffer 8, and Eosin 515 Trichrome, followed by dehydration through ethanol and clearing in xylene. E-cadherin and MUC2 slides were HIER (heat induced epitope retrieval) in pH 9.0 Tris-EDTA solution (10 mM Tris Base, 1 mM EDTA, 0.05% tween 20) for 3 minutes in Decloaking Chamber (Biocare Medical, Concord, CA). Slides were then cooled for 20 min, rinsed in 1x PBS, blocked with 10% goat serum (Sigma, St. Louis, MO) in PBS for 1 hour at room temperature, followed by primary antibody incubation in PBS with 1% goat serum and 0.1% Triton X-100 at 1:200 overnight at 4°C. The slides were then rinsed x 3 with 1x PBS and incubated with Alexafluor Goat anti-rabbit 488 at 1:1000 in PBS for 1 hr at room temperature. After the secondary incubation, slides were rinsed x 3 with 1x PBS followed by a DH_2_O rinse, and mounted with a cover slip using Prolong Gold with DAPI (Thermo Fisher Scientific, P36931). Images were acquired with a Nikon iLas2 TIRF/STORM/Epifluorescence microscope and mean pixel intensity of each stain was measured using ImageJ.

#### H&E histological analyses

Image acquisition and coding were performed by an independent investigator and all analyses (crypt length, number of goblet cells, and % area of inflammation) were conducted in a blinded manner. Crypt length and number of goblet cells per crypt were measured using QuPath software v0.6.0. Five crypts per mouse/image were analyzed, measuring the distance from the crypt base to the luminal surface and the number of goblet cells per crypt. The % area of inflammation, analyzing the same 5 crypts per mouse used for crypt length and number of goblet cells, was quantified using ImageJ. A uniform threshold was applied to the hematoxylin stain. Clustered cells were separated using the watershed function. Area of inflammation was quantified using the “Analyze Particles” function with a size threshold of 0-infinity pixel^2^ and circularity range of 0.00–1.00. Tumor numbers, grade, invasion and lymphovascular invasion events were scored according to published guidelines in a blinded manner^57,58^.

#### Fecal lipocalin-2 analysis

Fecal lipocalin-2 levels were quantified using a Mouse Lipocalin-2/NGAL DuoSet ELISA kit (R&D Systems, DY1857) according to the manufacturer’s instructions. High-binding 96-well plates (Corning, 3590) were coated overnight at room temperature with capture antibody diluted in PBS. Plates were washed with PBS containing 0.05% Tween-20 and blocked with 1% BSA in PBS for 1 h at room temperature. Fecal samples were homogenized in 300-500 μL PBS containing 0.1% Tween-20, centrifuged (18,200 x *g*, 10 min, 4°C), and the supernatant was collected for analysis. Supernatants were diluted 5x-25x in reagent diluent. Samples and standards were added to the plate and incubated for 2 h at room temperature. After washing, detection antibody and streptavidin-HRP were sequentially applied with intermediate washing steps. Signal was developed using TMB substrate (R&D Systems, DY999B) and stopped with stop solution (R&D Systems, DY994). Absorbance was measured at 450 nm with wavelength correction at 540 nm. Concentrations were calculated from the standard curve.

#### Serum IL-10 cytokine measurement

LEGENDplex™ Mouse Anti-Virus Response Panel with V-Bottom Plate (BioLegend, 740622) was used to measure serum IL-10 cytokine levels. For the standards, 25 µL of Matrix A was added into an assay plate, followed by adding 25 µL of serially diluted standard solutions. For serum samples, 25 µL of assay buffer was added into the assay plate, followed by adding 25 µL of 4-8x diluted serum samples. 25 µL of mixed beads were added into each well and the plate was shaken at 3,000 x *g* at room temperature for 2 h. After aspirating and washing the plate, 25 µL of detection antibodies were added into the plate followed by shaking at 3,000 x *g* at room temperature for 1 h. Then, 25 µL of SA-PE was added to each well, followed by shaking at 3,000 x *g* for 30 min at room temperature. After aspirating and washing the plate, the beads in the plate were resuspended in 150 µL of 1X wash buffer by pipetting and the plate was read on Attune NxT Flow Cytometer at the UW-Madison Carbone Cancer Center (UWCCC) Flow Cytometry Laboratory. Data were analyzed using the LEGENDplex™ Data Analysis Software (BioLegend). Bead populations were distinguished by FSC/SSC and internal dye intensity measured in the APC (670 nm) channel, and reporter signals were quantified in the PE (574 nm) channel.

### Statistical analysis

Data are presented as mean ± SEM for bar graphs and scatter dot plots, unless otherwise specified. Violin plots depict data distribution, with the central line indicating the median and dotted lines representing all data points. Each data point represents an individual biological replicate, unless otherwise specified, as defined in the figure legends. For comparisons between two groups, an unpaired two-tailed Welch’s t test was used. For non-normally distributed data, the Mann–Whitney test was applied. For comparisons among three or more groups, one-way ANOVA followed by appropriate post hoc multiple comparison tests was used. When normality assumptions were not met, the Kruskal–Wallis test was used. A power calculation was used to determine total number of animals required for each study. Briefly, a dichotomous endpoint for two independent study groups with an expected error in the mean of 10% and a difference of at least 75% was considered. A false positive rate of 10% and a false negative rate of 20% (power) identified a sample size of at least n=4 for each study group. A *p* value < 0.05 was considered statistically significant. Exact *p* values are reported in the figures.

## Graphics

The schematic illustrations were created using BioRender. Chemical structures were generated using ChemDraw. Molecular docking structures and structural visualizations were rendered using PyMOL.

## Data availability

This paper analyzes existing, publicly available human metagenomic datasets. Custom scripts are available on Github: https://github.com/allie-walker/MetagenomeGeneAbundance. The mass spectrometry proteomics data have been deposited to the ProteomeXchange Consortium with the data set identifier PXD081010. Proteomics data are publicly available as of the date of publication. Microbiome 16s rRNA sequencing data have been deposited in the NCBI Sequence Read Archive (SRA) database under accession number SRA: PRJNA1432080 (IL10KO vs. WT) and SRA: PRJNA1494587 (*EcN* administration experiments). Data are publicly available as of the date of publication. All other data reported in this paper will be shared by the lead contact upon request.

## Author contributions

Conceptualization and study design was carried out by R.D. and S.N.C. R.D. performed all *in vitro* and *in vivo* experiments along with tissue processing and data analysis with the help of J.C., Z.D., X.D., and K.M. Development of the folate metabolomics workflow and optimization of bacterial folate extraction and data processing was performed by J.W. and J.A.H. Z.D. generated CAPN2KO and CAPN2OE cell lines. C.Y. and S.N.C. performed CAPN2 bacterial expression, purification, and folate binding assays. S.N.C. generated *EcN glyA KO* under the guidance of C.L. Z.D. optimized and performed qPCR analysis of *EcN* colonization. J.G.V.V. performed target engagement proteomics. A.S.W. performed metagenomic analysis in human CRC datasets. R.D. and S.N.C. wrote the paper and prepared all figures. All authors edited and contributed to the critical review of the paper.

## Acknowledgements

This study was supported by the National Institutes of Health (R00 DK128503 to S.N.C., T32 GM135066 to J.W.), the Wisconsin Alumni Research Foundation, and the Department of Biochemistry at UW-Madison. We would like to thank the UWCC Experimental Animal Pathology Laboratory for all histology and immunofluorescent staining services, K.R. Weiss from the Biochemistry Optical Core at UW-Madison for assistance with fluorescence imaging, UWCC Flow Cytometry Laboratory, and C. Bingman at UW-Madison for the CAPN2 plasmid. We acknowledge the Lim and Simcox labs at UW-Madison for help with equipment. A special thank you to Drs. M. Alexander and D.A. Harris at UW-Madison for insightful guidance that strengthened this work. Finally, we are grateful to the members of the Chaudhari lab for technical assistance, helpful discussions, and critical reviews.

---

**Extended Data Fig. 1.**
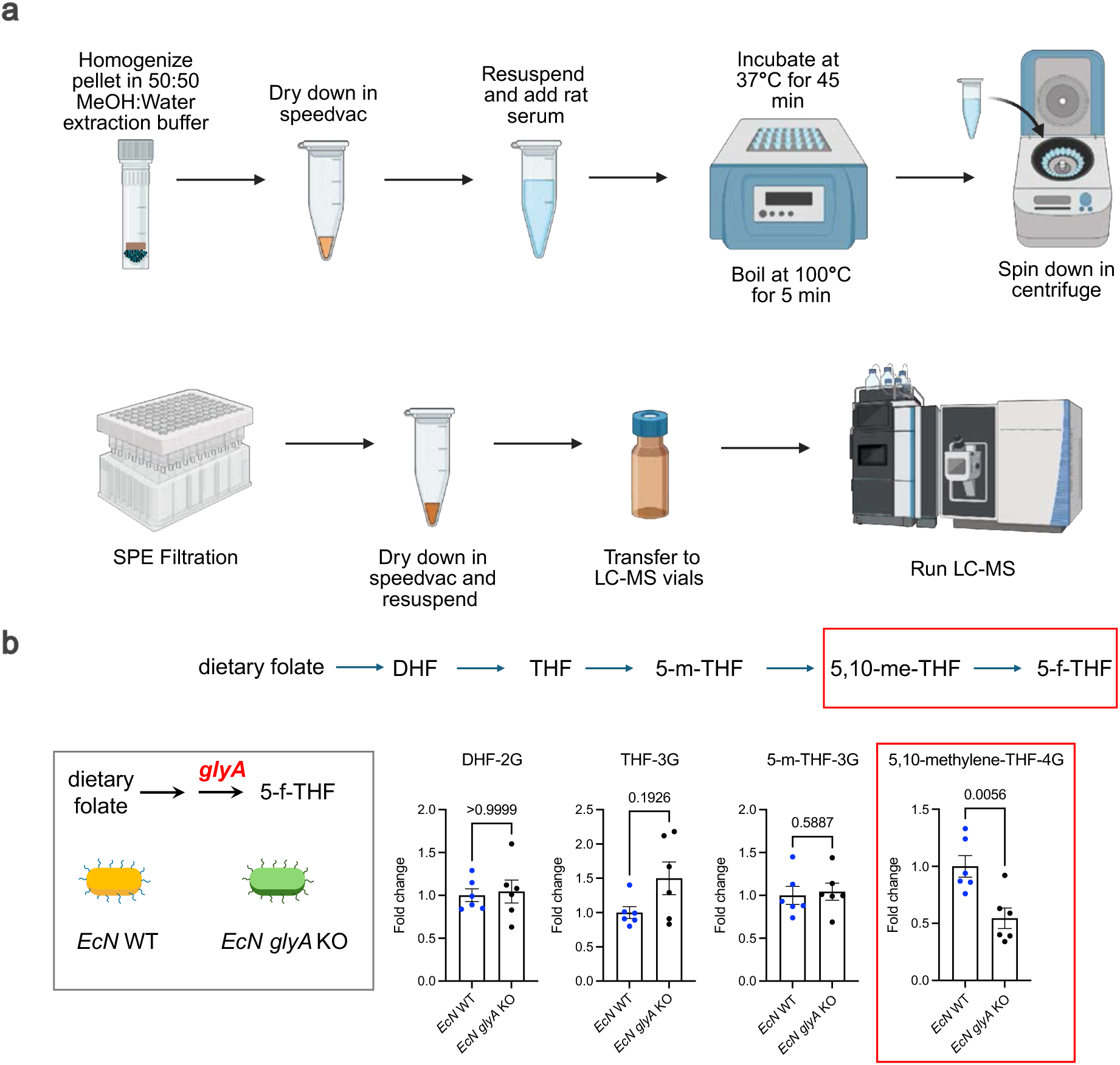
Genetic engineering of *EcN* to reduce folinic acid (5-f-THF) production. **a.** Schematic for LC-MS metabolomics for extraction and quantification of folate from bacterial cells. **b**. Schematic and quantification of dietary folate converted to dihydrofolate (DHF), tetrahydrofolate (THF), 5,10-methylene THF (5,10-me-THF) and 5-f-THF by *EcN* WT and *EcN glyA* KO. All statistically significant *p* values are indicated in each bar graph. Data not marked are not statistically significant (*p* > 0.05). All bar graphs are represented as mean ± SEM, data points represent biological replicates.

**Extended Data Fig. 2.**
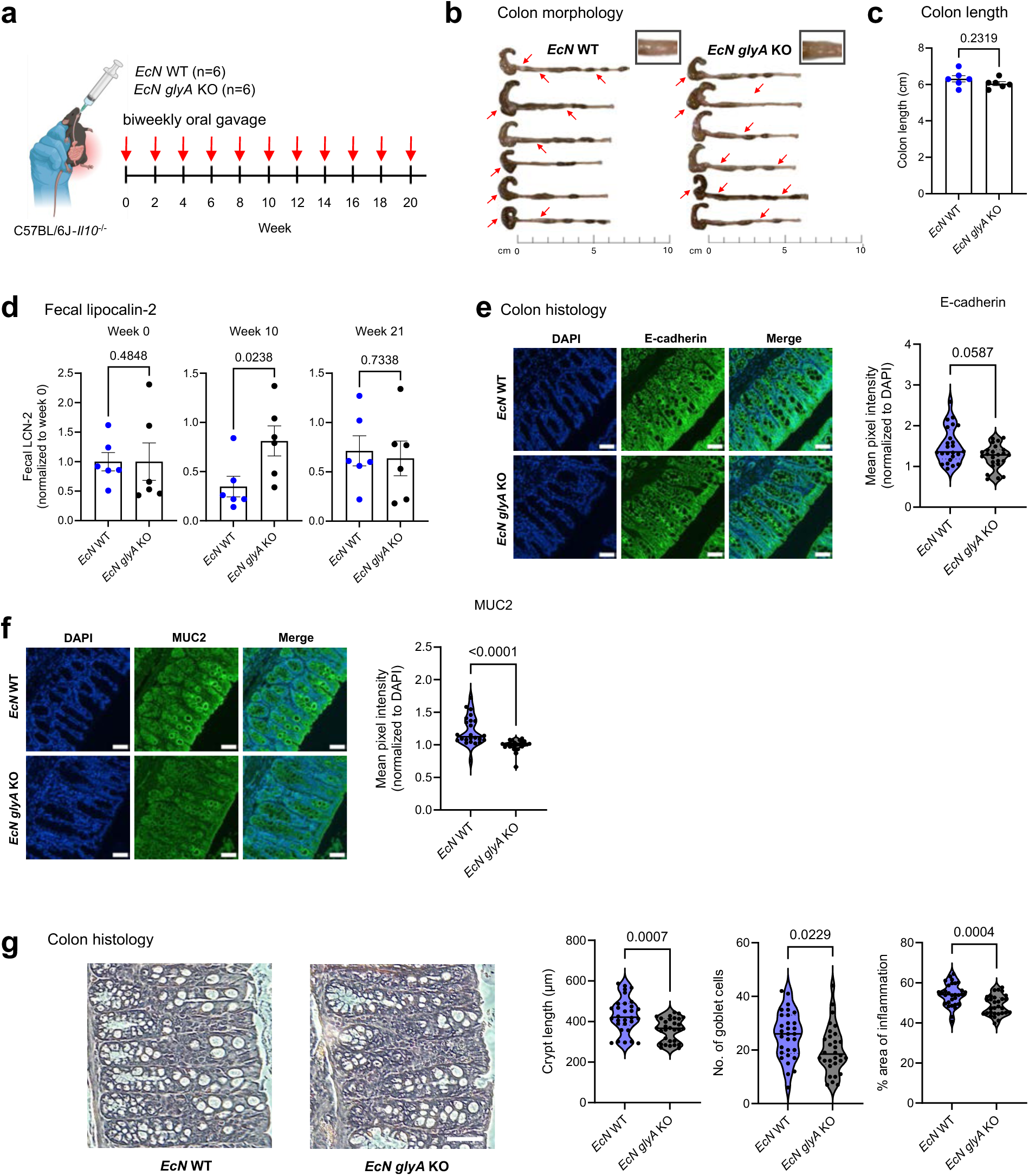
Microbial synthesis of folinic acid reduces CRC burden without mitigating inflammation. **a.** Schematic of a biweekly oral gavage of *E. coli* Nissle wild-type (WT) or *glyA* KO (10^9^ CFUs) administered to IL10KO mice at 11 weeks of age, followed by colon collection after 21 weeks. **b.** Colon images with irregularities marked with red arrows. **c.** Colon length (cm) of each mouse. (n=6 per group, two-tailed Welch’s t test) **d.** Fecal lipocalin-2 (LCN-2) levels measured by ELISA in samples collected at week 0, 10 and 21. Each n represents one mouse (n=6 per group, Mann-Whitney test). **e.** Representative images of mouse colons stained with E-cadherin and DAPI. Scale bar, 50 µm. Mean pixel intensity of E-cadherin was measured from 4 images per mouse colon using ImageJ and normalized to DAPI. (n=24 per group, Mann-Whitney test). **f.** Representative images of mouse colons stained with mucin 2 (MUC2) and DAPI. Scale bar, 50 µm. Mean pixel intensity of MUC2 was measured from 4 images per mouse colon using ImageJ and normalized to DAPI. (n=24 per group, Mann-Whitney test). **g.** Representative images of mouse colons stained with hematoxylin and eosin (H&E). Scale bar, 100 µm. All images were analyzed in a blinded manner. Crypt length (µm) and number of goblet cells were measured by QuPath and the % area of inflammation was measured by ImageJ based on 5 crypts per mouse. Each n represents one crypt. (n=30 per group; two-tailed Welch’s t test (number of goblet cells) and Mann-Whitney test (crypt length and % area of inflammation). All statistically significant *p* values are indicated in each bar graph. All bar graphs are represented as mean ± SEM, data points represent biological replicates.

**Extended Data Fig. 3.**
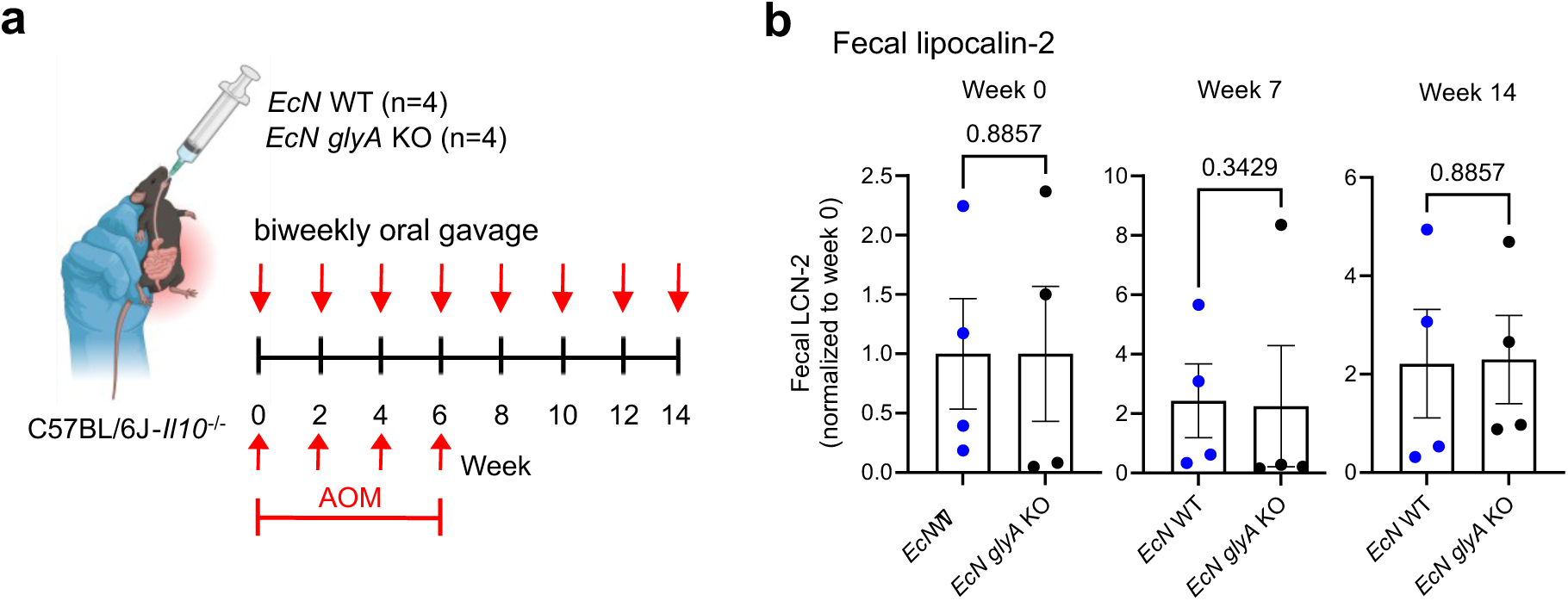
Microbial synthesis of folinic acid reduces CRC burden without mitigating inflammation. **a.** Schematic of weekly AOM treatment (10 mg/kg body weight) for six weeks and biweekly oral gavage of *E. coli* Nissle wild-type (WT) or *glyA* KO (10^9^ CFUs) administered to IL10KO mice at 16 weeks of age. **b.** Fecal lipocalin-2 (LCN-2) levels measured by ELISA in samples collected at week 0, 7 and 14. Each n represents one mouse (n=4 per group, two-tailed Welch’s t test). All *p* values are indicated in each graph. All bar graphs are represented as mean ± SEM, data points represent biological replicates.

## References

1. Piedbois, P. Modulation of fluorouracil by leucovorin in patients with advanced colorectal cancer: An updated meta-analysis. Journal of Clinical Oncology 22, 3766–3775 (2004).

2. Glimelius, B., Stintzing, S., Marshall, J., Yoshino, T. & de Gramont, A. Metastatic colorectal cancer: Advances in the folate-fluoropyrimidine chemotherapy backbone. Cancer Treat. Rev. 98, 102218 (2021).

3. Vodenkova, S. et al. 5-fluorouracil and other fluoropyrimidines in colorectal cancer: Past, present and future. Pharmacol. Ther. 206, 107447 (2020).

4. Zhang, N., Yin, Y., Xu, S. J. & Chen, W. S. 5-Fluorouracil: Mechanisms of Resistance and Reversal Strategies. Molecules 13, 1551 (2008).

5. Howard, S. C., McCormick, J., Pui, C.-H., Buddington, R. K. & Harvey, R. D. Preventing and Managing Toxicities of High-Dose Methotrexate. Oncologist 21, 1471–1482 (2016).

6. Menezo, Y., Elder, K., Clement, A. & Clement, P. Folic Acid, Folinic Acid, 5 Methyl TetraHydroFolate Supplementation for Mutations That Affect Epigenesis through the Folate and One-Carbon Cycles. Biomolecules 12, 197 (2022).

7. Wang, S. et al. Leucovorin Enhances the Anti-cancer Effect of Bortezomib in Colorectal Cancer Cells. Scientific Reports 2017 7:1 7, 682- (2017).

8. Colorectal cancer. https://www.who.int/news-room/fact-sheets/detail/colorectal-cancer.

9. Zalila-Kolsi, I., Dhieb, D., Osman, H. A. & Mekideche, H. The Gut Microbiota and Colorectal Cancer: Understanding the Link and Exploring Therapeutic Interventions. Biology (Basel). 14, 251 (2025).

10. Cao, Q., Yang, M. & Chen, M. Metabolic interactions: how gut microbial metabolites influence colorectal cancer. Front. Microbiol. 16, 1611698 (2025).

11. Liu, Y., Cheuk-Hay Lau, H. & Yu, J. Microbial metabolites in colorectal tumorigenesis and cancer therapy. Gut Microbes 15, (2023).

12. Wirbel, J. et al. Meta-analysis of fecal metagenomes reveals global microbial signatures that are specific for colorectal cancer. Nature Medicine 2019 25:4 25, 679–689 (2019).

13. Thomas, A. M. et al. Metagenomic analysis of colorectal cancer datasets identifies cross-cohort microbial diagnostic signatures and a link with choline degradation. Nature Medicine 2019 25:4 25, 667–678 (2019).

14. Dalal, N. et al. Gut microbiota-derived metabolites in CRC progression and causation. Journal of Cancer Research and Clinical Oncology 2021 147:11 147, 3141–3155 (2021).

15. Bautista, J. et al. Gut microbiome–driven colorectal cancer via immune, metabolic, neural, and endocrine axes reprogramming. npj Biofilms and Microbiomes 2026 12:1 12, 21-(2026).

16. Feitelson, M. A., Arzumanyan, A., Medhat, A. & Spector, I. Short-chain fatty acids in cancer pathogenesis. Cancer Metastasis Rev. 42, 677 (2023).

17. Williams, J. et al. Atlas of one-carbon metabolism in conventional and germ-free mice reveals folate as a key determinant of biochemical pathways. Nature Metabolism 2026 1–17 (2026) doi:10.1038/s42255-026-01489-w.

18. Misselbeck, K., Marchetti, L., Priami, C., Stover, P. J. & Field, M. S. The 5-formyltetrahydrofolate futile cycle reduces pathway stochasticity in an extended hybrid-stochastic model of folate-mediated one-carbon metabolism. Sci. Rep. 9, 4322 (2019).

19. Lin, J. M. G. et al. Metabolic modulation of transcription: The role of one-carbon metabolism. Cell Chem. Biol. 29, 1664–1679 (2022).

20. Jeanguenin, L. et al. Moonlighting Glutamate Formiminotransferases Can Functionally Replace 5-Formyltetrahydrofolate Cycloligase. Journal of Biological Chemistry 285, 41557–41566 (2010).

21. Berg, D. J. et al. Enterocolitis and colon cancer in interleukin-10-deficient mice are associated with aberrant cytokine production and CD4(+) TH1-like responses. Journal of Clinical Investigation 98, 1010 (1996).

22. Yang, Y. et al. Dysbiosis of human gut microbiome in young-onset colorectal cancer. Nature Communications 2021 12:1 12, 6757- (2021).

23. Yachida, S. et al. Metagenomic and metabolomic analyses reveal distinct stage-specific phenotypes of the gut microbiota in colorectal cancer. Nature Medicine 2019 25:6 25, 968–976 (2019).

24. Cheng, A. G. et al. Design, construction, and in vivo augmentation of a complex gut microbiome. Cell 185, 3617–3636.e19 (2022).

25. Van der Beek, J. N. et al. The effect of leucovorin rescue therapy on methotrexate-induced oral mucositis in the treatment of paediatric ALL: A systematic review. Crit. Rev. Oncol. Hematol. 142, 1–8 (2019).

26. Martínez-Maqueda, D., Miralles, B. & Recio, I. HT29 cell line. The Impact of Food Bioactives on Health: In Vitro and Ex Vivo Models 113–124 (2015) doi:10.1007/978-3-319-16104-4_11/SAVE-RESEARCH.

27. Vu, T. & Datta, P. K. Regulation of EMT in Colorectal Cancer: A Culprit in Metastasis. Cancers 2017, Vol. 9, Page 171 9, 171 (2017).

28. Hou, B. et al. LHPP suppresses colorectal cancer cell migration and invasion in vitro and in vivo by inhibiting Smad3 phosphorylation in the TGF-β pathway. Cell Death Discovery 2021 7:1 7, 273- (2021).

29. Fenouille, N. et al. Calpain 2-dependent IκBα degradation mediates CPT-11 secondary resistance in colorectal cancer xenografts. Journal of Pathology 227, 118–129 (2012).

30. Marciel, M. P. et al. Calpain-2 inhibitor treatment preferentially reduces tumor progression for human colon cancer cells expressing highest levels of this enzyme. Cancer Med. 7, 175–183 (2018).

31. Telechea-Fernández, M. et al. New localization and function of calpain-2 in nucleoli of colorectal cancer cells in ribosomal biogenesis: effect of KRAS status. Oncotarget 9, 9100 (2018).

32. Rose, A. H. et al. Calpain-2 inhibitor therapy reduces murine colitis and colitis associated cancer. Inflamm. Bowel Dis. 21, 2005 (2015).

33. Engevik, M. A. et al. Microbial Metabolic Capacity for Intestinal Folate Production and Modulation of Host Folate Receptors. Front. Microbiol. 10, (2019).

34. Datsenko, K. A. & Wanner, B. L. One-step inactivation of chromosomal genes in Escherichia coli K-12 using PCR products. Proc. Natl. Acad. Sci. U. S. A. 97, 6640–6645 (2000).

35. Brückner, M. et al. Detection and characterization of murine colitis and carcinogenesis by molecularly targeted contrast-enhanced ultrasound. World J. Gastroenterol. 23, 2899–2911 (2017).

36. Manicassamy, S., Prasad, P. D. & Swafford, D. Mouse Models of Colitis-Associated Colon Cancer. Methods in Molecular Biology 2224, 133–146 (2021).

37. Asaf, S. et al. Lipocalin 2—not only a biomarker: a study of current literature and systematic findings of ongoing clinical trials. Immunol. Res. 71, 287 (2022).

38. Krause, P. et al. IL-10-producing intestinal macrophages prevent excessive antibacterial innate immunity by limiting IL-23 synthesis. Nature Communications 2015 6:1 6, 7055-(2015).

39. Rothemich, A. & Arthur, J. C. The azoxymethane/Il10 −/− model of colitis-associated cancer (CAC). Methods in Molecular Biology 1960, 215–225 (2019).

40. Wang, K. et al. Gut dysfunction may be the source of pathological aggregation of alpha-synuclein in the central nervous system through Paraquat exposure in mice. Ecotoxicol. Environ. Saf. 246, 114152 (2022).

41. Mini, E., Trave, F., Rustum, Y. M. & Bertino, J. R. Enhancement of the antitumor effects of 5-fluorouracil by folinic acid. Pharmacol. Ther. 47, 1–19 (1990).

42. Jonker, D., Rumble, R., Maroun, J. & Care, the G. C. D. S. G. C. C. O. P. in E.-B. Role of oxaliplatin combined with 5-fluorouracil and folinic acid in the first- and second-line treatment of advanced colorectal cancer. Current Oncology 13, 173 (2006).

43. Soveri, L. M. et al. Association of adverse events and survival in colorectal cancer patients treated with adjuvant 5-fluorouracil and leucovorin: Is efficacy an impact of toxicity? Eur. J. Cancer 50, 2966–2974 (2014).

44. Peng, X., Yang, R., Song, J., Wang, X. & Dong, W. Calpain2 Upregulation Regulates EMT-Mediated Pancreatic Cancer Metastasis via the Wnt/β-Catenin Signaling Pathway. Front. Med. (Lausanne). 9, 783592 (2022).

45. Hara, H., Goshima, H., Hoshino, D. & Honda, M. Loss of Calpain-2 attenuates TGF-β–driven Epithelial–Mesenchymal transition and invasive behavior in A549 lung cancer cells. Biochem. Biophys. Res. Commun. 790, (2025).

46. Harper, D. et al. Calpain-1 and Calpain-2 Promote Breast Cancer Metastasis. Cells 14, (2025).

47. Zhang, G. et al. Calpain 2 knockdown promotes cell apoptosis and restores gefitinib sensitivity through epidermal growth factor receptor/protein kinase B/survivin signaling. Oncol. Rep. 40, 1937 (2018).

48. Iyer, R. & Tomar, S. K. Folate: a functional food constituent. J. Food Sci. 74, (2009).

49. Li, T. et al. Folate exposures and risk of colorectal cancer: an umbrella review of meta-analyses of observational studies and randomised controlled trials. BMJ Open 15, e103637 (2025).

50. Callahan, B. J. et al. DADA2: High-resolution sample inference from Illumina amplicon data. Nature Methods 2016 13:7 13, 581–583 (2016).

51. Caporaso, J. G. et al. QIIME allows analysis of high-throughput community sequencing data. Nature Methods 2010 7:5 7, 335–336 (2010).

52. Hughes, C. S. et al. Single-pot, solid-phase-enhanced sample preparation for proteomics experiments. Nat. Protoc. 14, 68–85 (2019).

53. Li, J. et al. TMTpro reagents: a set of isobaric labeling mass tags enables simultaneous proteome-wide measurements across 16 samples. Nature Methods 2020 17:4 17, 399–404 (2020).

54. Thompson, A. et al. TMTpro: Design, Synthesis, and Initial Evaluation of a Proline-Based Isobaric 16-Plex Tandem Mass Tag Reagent Set. Anal. Chem. 91, 15941–15950 (2019).

55. Huttlin, E. L. et al. A tissue-specific atlas of mouse protein phosphorylation and expression. Cell 143, 1174–1189 (2010).

56. Savitski, M. M., Wilhelm, M., Hahne, H., Kuster, B. & Bantscheff, M. A scalable approach for protein false discovery rate estimation in large proteomic data sets. Molecular and Cellular Proteomics 14, 2394–2404 (2015).

57. Fleming, M., Ravula, S., Tatishchev, S. F. & Wang, H. L. Colorectal carcinoma: Pathologic aspects. J. Gastrointest. Oncol. 3, 153–173 (2012).

58. Kim, B. H. et al. Standardized Pathology Report for Colorectal Cancer, 2nd Edition. J. Pathol. Transl. Med. 54, 1–19 (2019).

